# Unbiased complexome profiling and global proteomics analysis reveals mitochondrial impairment and potential changes at the intercalated disk in presymptomatic R14^Δ/+^ mice hearts

**DOI:** 10.1101/2024.03.21.586093

**Authors:** Brian Foo, Hugo Amedei, Surmeet Kaur, Samir Jaawan, Angela Boshnakovska, Tanja Gall, Rudolf A. de Boer, Herman H.W. Silljé, Henning Urlaub, Peter Rehling, Christof Lenz, Stephan E. Lehnart

## Abstract

**Background:** Phospholamban (PLN) is a sarco-endoplasmic reticulum (SER) membrane protein that regulates cardiac contraction/relaxation by reversibly inhibiting the SERCA2a Ca^2+^-reuptake pump. The R14Δ-PLN mutation causes severe cardiomyopathy that is resistant to conventional treatment. Protein complexes and higher-order supercomplexes such as intercalated disk components and Ca^+2^-cycling domains underlie many critical cardiac functions, a subset of which may be disrupted by R14Δ-PLN.

**Methods:** We developed an improved complexome profiling (CP) workflow specifically geared towards identifying disruption of very high molecular-weight (>2 MDa) protein complexes and supercomplexes in presymptomatic R14^Δ/+^ mice hearts. Ventricular tissues were homogenized under non-denaturing conditions, fractionated by size-exclusion chromatography (SEC) and subjected to quantitative data-independent acquisition mass spectrometry (DIA-MS) proteomics analysis. Systematic analysis of CP data using conventional strategies yielded limited insights, likely due to underrepresentation of cardiac-specific complexes in the curated protein complex databases used as ground-truth for analysis. We thus developed PERCOM: a novel data analysis strategy that does not rely upon protein complex databases and can, furthermore, be implemented on widely available spreadsheet software.

**Results:** SEC-DIA-MS coupled with PERCOM identified 296 proteins with disrupted elution profiles in presymptomatic 28wk-old R14^Δ/+^ mice. Hits were significantly enriched for mitochondrial and intercalated disk (ICD) components. Alterations to mitochondrial and ICD supercomplexes were observed in mice as young as 9wks of age and were associated with reduced expression of mitochondrial proteins and maximal oxygen consumption rate.

**Conclusion:** Using a novel CP workflow, we identify mitochondrial alterations as an early-stage R14Δ-PLN event and provide preliminary data showing effects at the ICD. These molecular components underlie critical cardiac functions and their alteration at a young age may contribute to R14Δ-PLN pathogenesis.

## Introduction

Cardiac contraction/relaxation is regulated by controlled cycles of Ca^2+^ release and reuptake from the sarco-endoplasmic reticulum (SER), mediated by the Ryanodine receptor 2 (RyR2) Ca^2+^-release channel and the sarcoplasmic/endoplasmic reticulum (SER) Ca^2+^ ATPase 2a (SERCA2a) reuptake pump, respectively^1^. Phospholamban (PLN) is a 52-aa tail-anchored SER-membrane protein that plays a key role in regulating SERCA2a function^2^. PLN reversibly binds-to and inhibits SERCA2a; this interaction is negatively regulated by phosphorylation of PLN at Ser16 by protein kinase A (PKA, in response to adrenergic signaling) and Thr17 by Ca^2+^/calmodulin-dependent protein kinase II (CaMKII, in response to adrenergic signaling and increased cytosolic Ca^2+^)^2^. Several definitively pathogenic disease-associated PLN mutations have been described to-date, including R9C^3^, L39stop^4^ and R14Δ^5^.

The R14Δ mutation is found worldwide, with a particularly high prevalence the Netherlands and Greece where it is among the leading forms of genetic cardiomyopathy^6^. It is associated with arrhythmogenic and/or dilated cardiomyopathy (ACM/DCM), frequently resulting in heart failure at adolescence and beyond that is resistant to conventional treatment^7,8^. On the molecular level, R14Δ-PLN is associated with loss of PKA binding, resulting in PLN hypo-phosphorylation leading to altered SERCA2a regulation and Ca^2+^-handling dysfunction^5,9,10^. On the cellular level, R14Δ-PLN is associated with a complex phenotype involving accelerated Ca^2+^-handling dynamics, formation of PLN-positive perinuclear SER clusters, upregulated UPR signaling and mitochondrial dysregulation^7,11–14^. Despite over a decade of work, the mechanisms linking R14Δ-PLN to disease pathology remain poorly understood.

Many proteins assemble into multimeric and/or multi-protein complexes with well-defined molecular composition and function^15,16^. Notable examples include mitochondrial respiratory-chain complexes I-V^17^, SERCA2a and Na^+^/K^+^-ATPase dimers^18,19^ and voltage-gated K^+^-channel tetramers^20^. Protein complexes may form further higher-order supercomplexes and subcellular nanodomains, which may be characterized by larger size (> 1MDa), fluid composition and/or stoichiometry, and coordination of several distinct functions such as signaling and membrane-tethering^15,21^. There is a growing appreciation that disruption of these assemblies may contribute to cardiac diseases. Examples include disruption of the SER Ca^2+^-handling supercomplex in atrial fibrillation^22^, SER-plasma membrane (PM) Ca^2+^-release unit (CRU) in heart failure^23^, and intercalated disk components during ischemia-reperfusion injury^24^ and ACM^25^.

Complexome Profiling (CP) is a systems-biology approach to study changes in the composition of protein complexes. Data acquisition and analysis workflows are summarized in Figure 1a. Briefly: biological samples are homogenized under non-denaturing conditions, fractionated based on molecular weight (MW) using techniques such as gradient centrifugation^26,27^, blue-native electrophoresis^28^ (BNE) or size-exclusion chromatography^29–31^ (SEC), and the resultant fractions analyzed by bottom-up mass spectrometry (MS)-based proteomics. Raw data output may consist of a matrix containing the abundance of each detected protein across each elution fraction (with elution fraction number correlating with MW). Any protein may be expected to elute at the MW-fraction corresponding to its predicted MW; in the case of PLN at a predicted MW of ∼6 kDa. Should the protein be incorporated into higher-MW protein complexes or supercomplexes, a subpopulation would also elute at higher-MW fractions corresponding to the apparent MW of that complex; in the case of PLN incorporated into the SER Ca^+2^-handling supercomplex, we observe a broad peak at 2-5 Mda (Figure 2a-c)^22^. Furthermore, proteins that are part of the same complex/supercomplex would be expected to at-least partially co-elute at these high-MW fractions.

**Figure 1:**
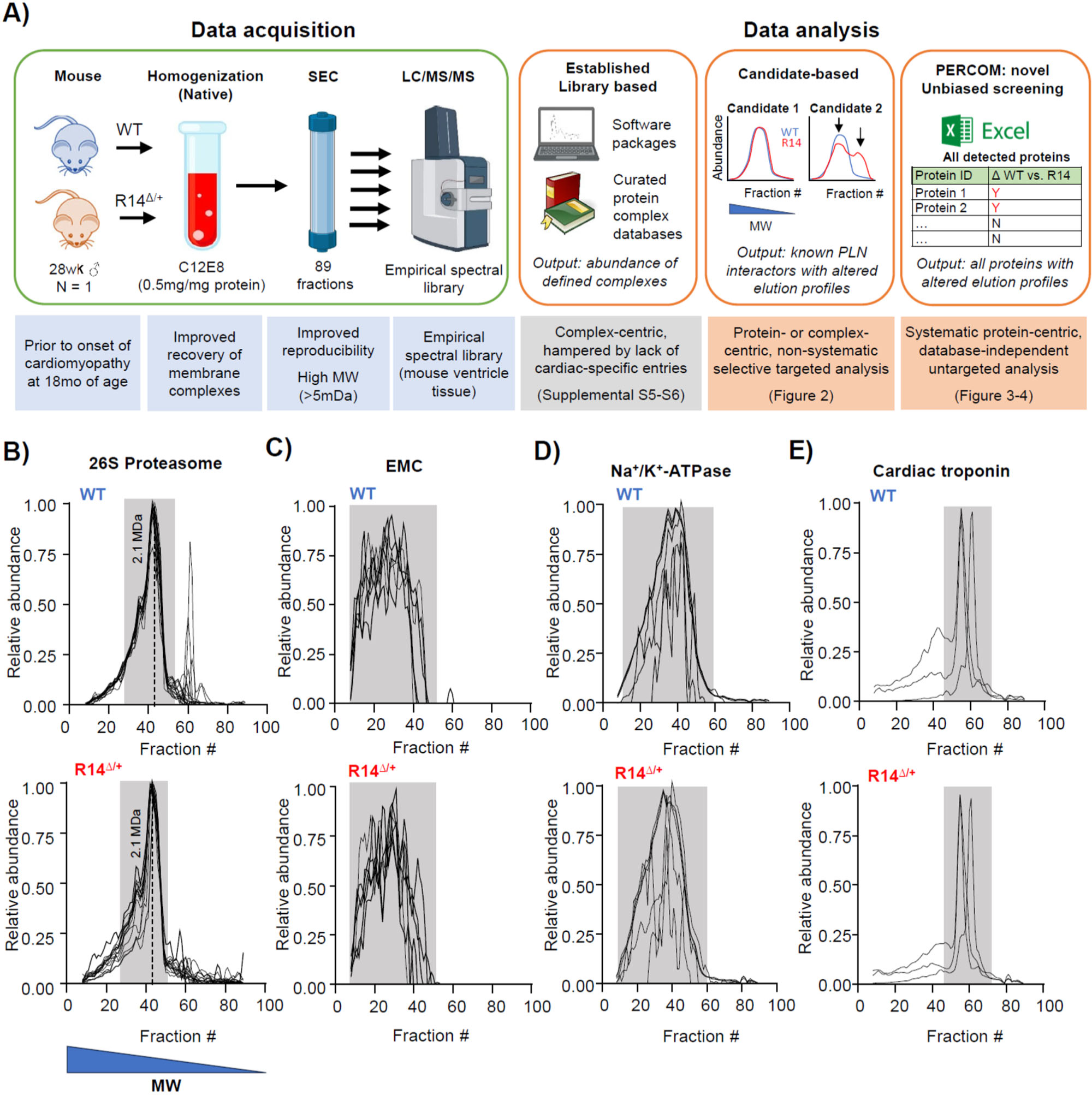
Development of a high MW complexome profiling workflow. **A)** SEC-MS based complexome profiling workflow for cardiac samples. Data-acquisition: Left-ventricular (LV) tissue was dissected from sibling-matched presymptomatic 28-week-old ♂ WT and R14^Δ/+^ mice (N = 1). Membrane fractions were enriched by differential centrifugation, solubilized in C12E8 (0.5 mg/mg protein) and separated into 89 fractions by SEC. Homogenization and fractionation workflows were optimized for preservation of membrane protein complexes, detection at high-MW ranges and improved reproducibility (see text). Each fraction was then subject to MS-based proteomics analysis using a dedicated empirical spectral library for mouse ventricular tissue for ground-truth. Data-analysis: three modes of data-analysis were employed: a traditional complex-centric automated approach comparing the abundance of known protein complexes (defined within the CORUM 4.0 protein complex database) between samples (Supplemental figure S6), a candidate-based approach based upon manual comparison of known PLN-interacting proteins in WT and R14^Δ/+^ mice (Figure 2), and PERCOM: a novel unbiased protein-centric screening strategy implemented on Microsoft Excel which systematically analyzes the elution profiles of all detected proteins and outputs a list of proteins with altered elution profiles (Figure 3). **B-E)** SEC-MS based data-acquisition workflow preserves protein complexes from diverse subcellular compartments. Constituent proteins of the 26S proteasome (soluble cytosolic complex), ER membrane complex (EMC, integral SER membrane), Na^+^/K^+^-ATPase (integral plasma membrane) and cardiac troponin (sarcomere) were detected from WT (top) and R14^Δ/+^ (bottom) mouse tissue. Elution profiles represented as a plot of relative abundance vs. fraction #, with each trace representing the profile from a single protein (protein IDs listed in Table 2). Higher fraction # corresponds to lower MW (panel B, bottom). Dotted line: 2.1 MDa MW marker (Full MW calibration in Supplemental Figure S3). Peaks corresponding to each complex manually evaluated (grey underlay). Fractions 1-7 were not analyzed due to lack of protein as detected by UV spectroscopy (Supplemental Figure S3). Elution profiles subject to curve smoothing (4th order polynomial, 2nd neighbor). Points lying below the horizontal axis resulting from smoothing removed for clarity.

**Figure 2:**
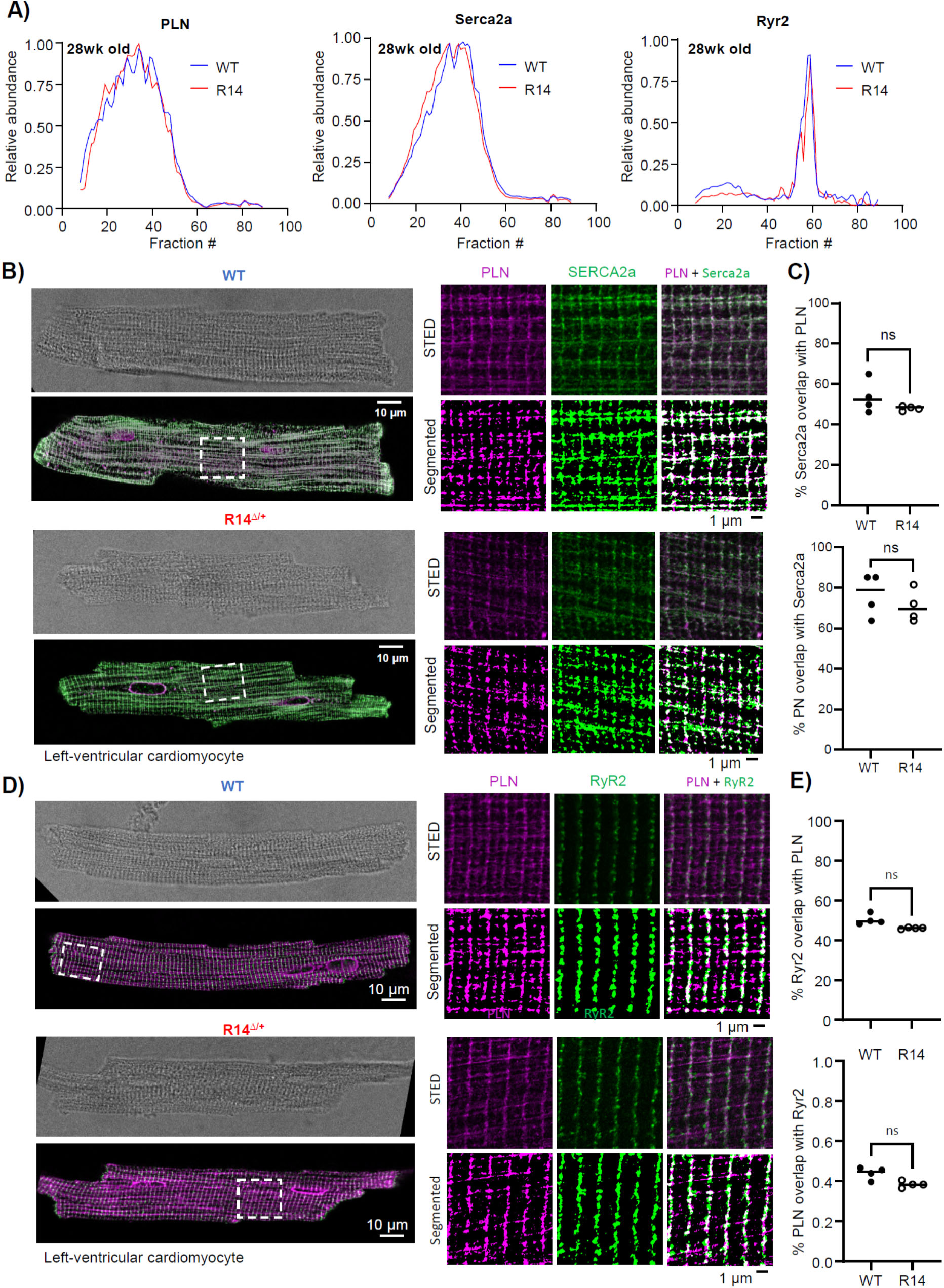
The PLN-containing SER Ca^2+^-handling supercomplex is not affected in adult 21-28wk-old R14^Δ/+^ mice. **A)** Elution profiles of PLN and associated proteins SERC2a and RyR2 from 28 wk-old ♂ WT and R14^Δ/+^ mouse tissue. Experimental workflow shown in Figure 1a (N = 1). Elution profiles subject to curve smoothing as in Figure 1b-e. **B-C)** PLN colocalization with SERCA2a evaluated by STED super-resolution microscopy in 21-28 wk-old ♂ mice (N = 4). Fraction of PLN pixels overlapping with SERC2a, and fraction of SERCA2a pixels overlapping with PLN quantified. **D-E)** PLN colocalization with Ryr2 evaluated by STED super-resolution microscopy as in panels b-c. Statistical significance determined using unpaired T-test with Bonferroni correction for multiple comparison (ns = no significant difference).

State-of-the-art approaches for systematic analysis of CP data utilize software packages or scripts such as CCprofiler^32,33^ or ComplexBrowser^34^ to detect and quantify the abundance of known protein complexes. Protein complex database such as CORUM^35^ or ComplexPortal^36^ provide ground-truth for these targeted “complex-centric” detection/quantification strategies. Alternatively, the fraction profiles for candidate proteins of interest may be manually compared.

While these techniques have been used with great success to identify co-migrating/eluting proteins in immortalized cell lines and *in-vitro* systems, allowing for detailed characterization of known protein complexes and *de novo* identification of new complexes^32,33,37,38^, several shortcomings in both data acquisition and data analysis steps prevent widespread application of this tool to identify disease-associated changes in specialized tissue types such as the myocardium. Successful data acquisition requires optimized workflows for homogenization and fractionation of complex tissues under native conditions. With regards to data analysis: existing protein complex databases used as “ground truth” are curated predominantly from immortalized and/or cancerous cell lines. As a consequence, cardiac-specific complexes such as CRUs or ICD components are severely underrepresented (Supplemental S5), inhibiting their use in cardiac research. In addition, most of the available data analysis workflows require scripting expertise to implement, limiting their accessibility.

Here, we developed a SEC-DIA-MS CP data acquisition strategy specifically geared towards fractionation of very high apparent MW protein complexes and supercomplexes (for example, the aforementioned SER Ca^2+^-handling supercomplex at ∼5 MDa^22^) from cardiac tissue. This was coupled to PERCOM, a novel untargeted data analysis workflow. PERCOM is a “protein-centric” approach that identifies individual proteins with altered elution profiles between two different conditions and is thus independent of curated protein complex databases. In addition, it can be implemented on common spreadsheet software such as Microsoft Excel without scripting expertise. SEC-DIA-MS/PERCOM analysis was performed on 28wk-old wild-type (WT) and heterozygous R14^Δ/+^ mice^7^. These animals, while exhibiting a mild cardiac conduction defect^7,39^, lack the severe cardiomyopathy, fibrosis and perinuclear cluster formation observed in end-stage human-patient hearts and thus reflect an early (presymptomatic) disease state^7,11^. Out of 3055 detected proteins, we identified 296 with altered elution profiles in R14^Δ/+^ mice. Gene ontology analysis revealed a disproportionate number of proteins involved in mitochondrial membrane organization, oxidative phosphorylation and intercalated disk components important for maintaining intercellular conduction and connectivity. Alterations in mitochondrial supercomplexes was associated with impaired mitochondrial function and reduced mitochondrial protein abundance. Lastly, these alterations were confirmed in juvenile (9wk-old) R14^Δ/+^ mice, suggestive of very-early disease events.

## Results

### 1. Development of a SEC-DIA-MS data acquisition workflow and application to cardiac tissue samples

Fractionation and data acquisition workflows for CP of cardiac samples frequently involve solubilization of either mitochondrial or membrane fractions in digitonin, followed by BNE-based fractionation and MS-based protein detection^22,40–42^. We improved upon this established workflow via the use of SEC-based fractionation techniques in lieu of BNE, and selection of an alternate detergent to improve preservation of membrane protein complexes. SEC-based fractionation offers several benefits over BNE such as the potential for enhanced resolution at very high MW-ranges (up to ∼7.5 MDa in our system, Supplemental Figure S3), enhanced reproducibility, increased loadability, and the potential to retain protein complexes in their solubilized native/active state to facilitate additional downstream analysis such as enzyme activity assays or protein/protein crosslinking mass spectrometry (XL-MS)^43^. In particular, enhanced reproducibility is crucial in comparing elution profiles between WT and disease conditions as in this work, and the increased MW-range will facilitate the detection of higher-order assemblies such as the ∼5 MDa PLN-containing SER Ca^2+^-handling supercomplex^22^.

Here, we performed SEC-CP on presymptomatic sibling-matched 28wk-old male WT and R14^Δ/+^ mice (N = 1). Left-ventricular (LV)-tissues were homogenized and solubilized in octaethylene glycol monododecyl ether (C12E8, 0.5mg per mg protein)^44^. C12E8 is a non-ionic detergent mostly used in the biophysical characterization of proteins in membrane bilayers^44,45^ and has recently found application in the analysis of protein complexes under near-native conditions by mass spectrometry^46,47^. We found this detergent to best preserve the integrity of membrane protein complexes in cardiac samples (Supplemental Figures S1-2). Following solubilization, samples were fractionated into 89 fractions using a novel 1000 Å pore size SEC column chemistry that has recently become commercially available and enables separation of biomolecular assemblies up to 7.5 MDa apparent MW ^48^ (Bio-SEC-5, 5 µm, 1000 Å; 300×7.8 mm, Agilent, USA, Supplemental Figure S3). The 82 protein-containing fractions were analyzed by bottom-up data-independent acquisition mass spectrometry (DIA-MS) using a custom annotated MS/MS spectral library to obtain global proteome profiles across the apparent MW dimension (Figure 1a, Supplemental Spreadsheets T1-2).

To our knowledge, this is the first successful application of SEC-DIA-MS sample-preparation workflows to myocardial tissue. As a first benchmark, we looked for the 26S proteasome which is a well-defined ∼2 MDa supercomplex. Our workflow was able to resolve a tight peak of 26S proteasomal proteins at the correct MW, validating our sample preparation workflow (Figure 1b). We were also able to resolve many other established protein complexes located in the cytosol (COP9 signalosome), plasma-membrane (Na^+^/K^+^-ATPase, Cav1.2 and Kir6.2 multimers), SER (EMC complex, Sec61) and sarcomere (troponin, titin) (Figure 1c-e, Supplemental Figure S3). Importantly, we did not observe notable profile changes between our WT and R14^Δ/+^ tissue samples, confirming the reproducibility of our workflow.

### 2. Candidate-based analysis shows no change in the PLN-containing SER Ca^2+^-handling supercomplex

We previously employed BNE-based complexome profiling of digitonin-solubilized mouse-heart and human-atrial tissues to identify a high-MW (∼5 MDa) SER Ca^2+^-handling supercomplex containing PLN, SERCA2a, RyR2, JPH2 (Junctophilin-2, a Cav1.2/RyR2 and SER tethering protein) and PP1R3A (a protein phosphatase 1 adaptor protein)^22^. This PLN-containing supercomplex, with a clear potential mechanistic link to the R14Δ-PLN Ca^2+^-cycling defect, is an obvious candidate for interrogation. We were able to robustly detect the SER Ca^2+^-handling supercomplex via SEC-MS complexome profiling; however, no notable difference was found between the elution profiles of PLN, SERCA2a and RyR2 from WT and R14^Δ/+^ mice (Figure 2a-c)^55^. Orthogonal evaluation was obtained via STED super-resolution microscopy of PLN-SERCA2a and PLN-RyR2 colocalization. Consistent with our previous findings^22^, significant PLN/SERCA2a and PLN/RyR2 overlap was observed; however colocalization was not significantly altered in R14^Δ/+^ mice (Figure 2b-e, Supplemental Figure S4). Recent work suggests that the R14Δ mutation alters the interaction between PLN and known binding partners such as HAX-1, HRC and SERCA2a^9^. In addition, an increased stability of the PLN-R14Δ pentamer has also been reported in HEK293 cells overexpressing PLN^49^ . While these prior findings indicate changes at the level of individual binding interactions, these do not appear to translate into observable disruption of the functional supercomplex.

### 3. PERCOM: an accessible and unbiased protein-centric data analysis workflow

The complex cellular phenotype associated with R14Δ-PLN, which includes impaired ER-mitochondrial contact^50^, mitochondrial dysregulation^14^, formation of PLN-positive perinuclear clusters^7,12^ and UPR signalling^13^ suggests alterations in other protein complexes/supercomplexes that may be functionally and/or spatially remote from the PLN-containing SER-Ca^2+^-handling supercomplex. Systematic analysis of CP data generally involves the use of software packages (ex. CCprofiler^32^ or ComplexBrowser^34^) that evaluate the presence and abundance of known annotated protein complexes based on the coelution of known subunits, with ground truth provided by external annotated protein databases such as ComplexPortal^51^ or CORUM^35^. This approach has been used with success to identify changes in well-established protein complexes, such as the respiratory chain complexes I-V in disease states^52^; however, several shortcomings limit their application to this study. i) Protein complex databases document well-defined and established complexes such as the T-cell receptor complex^53^, K^+^-channel tetramers^20^ and the Cop9 signalosome^54^; however, higher-order supercomplexes, which may have poorly-defined and fluid stoichiometry and/or composition (such as the aforementioned SER Ca^2+^-handling supercomplex^22^, intercalated disks components^55^ and signaling microdomains^56^), may not be represented. ii) These databases largely represent data from non-cardiac cell lines. For example: less than 1% of CORUM 4.0 entries are derived from primary cardiac material (Supplemental S5). As a consequence, many key cardiac-specific proteins (such as JPH2) and supercomplexes (such as the SER Ca^2+^-handling supercomplex^22^) are absent from both the ComplexPortal and CORUM databases. iii) More generally, these databases represent only a subset of the human proteome (22% for CORUM 4.0^35^). Taken together, traditional targeted “complex-centric” analysis strategies may be restrictive to a narrow subset of well-established protein complexes curated from non-cardiac cells, and thus fail to detect changes in cardiac-specific complexes and/or less well-defined higher-order supercomplexes. Lastly, these packages frequently require computer programming expertise to implement, which may present a barrier to some groups.

To illustrate these limitations, we attempted to apply traditional database-dependent data analysis workflows to our cardiac CP data. We employed a custom R-software package to identify and quantify, within our CP dataset, the abundance of protein complexes defined by the CORUM 4.0 protein complex database^35^. This workflow was able to identify and quantify the abundance of 57 annotated protein complexes in both WT and R14^Δ/+^ cardiac samples (Supplemental Figure S6). Nonetheless, the majority of detected complexes were not cardiac specific and, vice-versa, several noteworthy cardiac-specific complexes such as intercalated disk components and the SER Ca^2+^-handling supercomplex were not identified despite the constituent components co-eluting within our dataset (Figure 2a-c and Figure 9). Lastly, higher-order supercomplexes, such as the well-characterized respiratory supercomplex (RSC) containing mitochondrial complexes CI, III and IV^57^, were not detected, despite strong co-elution and detection of the individual constituent complexes (in the case of the RSC: mitochondrial complexes CI, III and IV, Figure 5 and Supplemental Figure S6).

To overcome these limitations, we developed PERCOM, a novel “protein-centric” analysis workflow with two main objectives: 1) to enable identification of individual proteins with altered elution profiles (as opposed to established and annotated protein complexes) in order to avoid the limitations of protein complex databases, and 2) to be implemented using common data processing software (ex. Microsoft Excel, Figure 1a). The data analysis workflow is summarized in Figure 3a. Input data consist of one matrix of protein abundances (either raw or relative intensities) per condition (in this case: a signle WT and R14^Δ/+^ mouse, N = 1). Protein ID and fraction number are organized by row and column, respectively. The elution profiles of each protein are then compared based on two parameters: Pearson’s coefficient of correlation (R-value) and the center-of-mass (CoM), giving rise to the name of this analysis: PERCOM. Both CoM and Pearson’s score can be visualized on a protein elution plot of protein abundance vs. fraction number (Figure 3b-c). CoM is defined as the fraction which evenly divides the area-under-the-curve (50% on either side, Figure 3b). The change in CoM (ΔCoM) represents the difference in CoM between the WT and R14^Δ/+^. The Pearson’s coefficient is calculated using the typical formula (Figure 3c). The combination of ΔCoM and Pearson’s coefficient was chosen to maximize detection of proteins with altered distribution profiles in a complementary manner. We postulate that ΔCoM would detect proteins with elution peaks slightly shifted or skewed towards lower-MW fractions, which may indicate subtle changes in supercomplex composition and/or assembly efficiency (Figure 3d-e). In contrast, Pearson’s coefficient would detect proteins with significant changes in elution profile shape, which may be consistent with complete disruption or reorganization of a supercomplex (Figure 3f).

**Figure 3:**
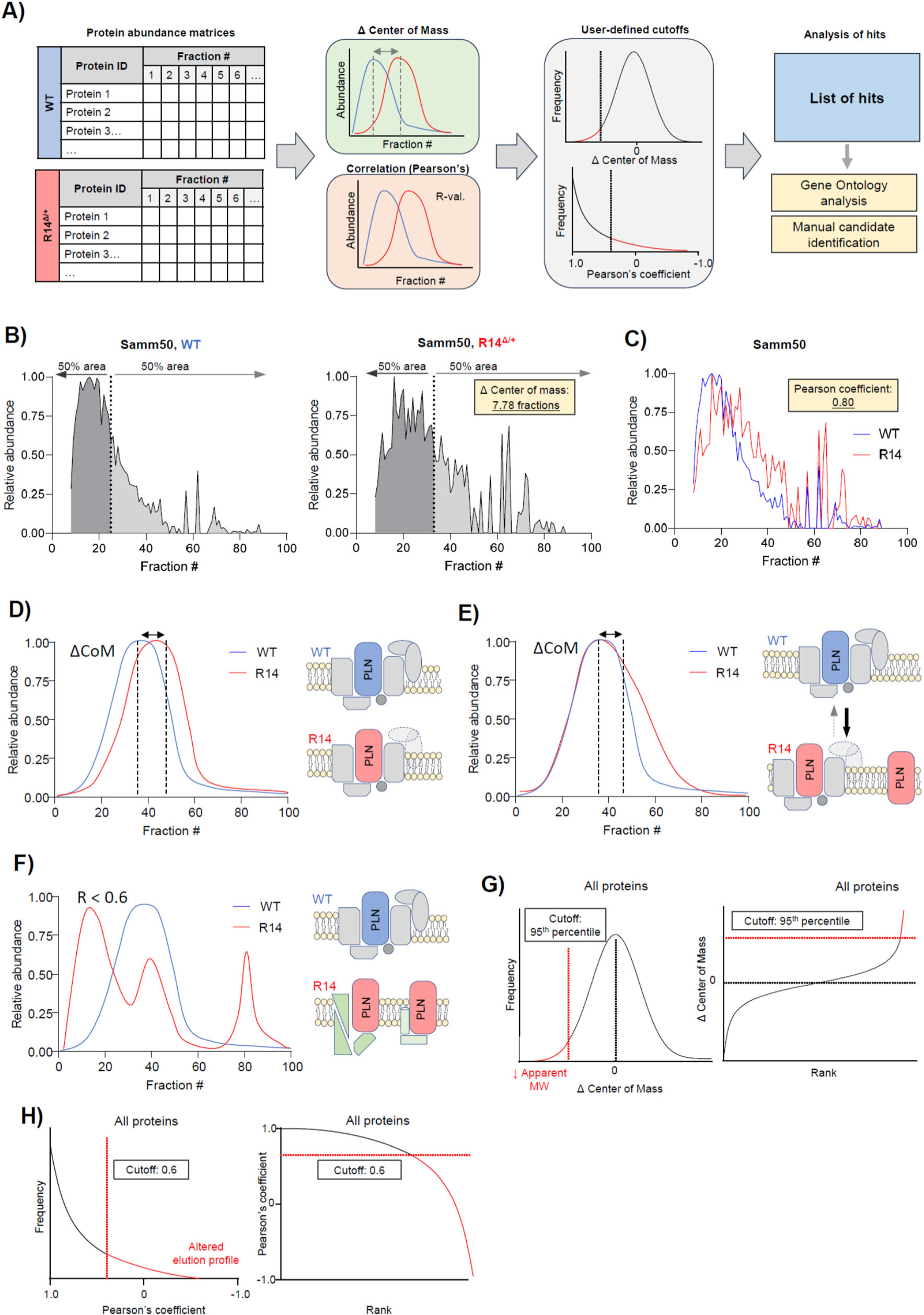
PERCOM, an unbiased and easily-accessible protein-centric analysis workflow. **A)** Graphical illustration of PERCOM data analysis strategy. Input data formatted as a matrix of protein abundances where protein ID and fraction number are organized by row and column, respectively, and each cell contains a value representing protein abundance. Each condition is represented by an individual matrix: in this case, WT and R14^Δ/+^. Two parameters are analyzed: the change (Δ) of center of mass (CoM) and Pearson’s correlation coefficient (R-value). Cutoffs are then applied to identify hits, which may then be subject to candidate analysis of individual proteins or interest or gene-ontology analysis to identify cellular components and/or biological pathways overrepresented within hits. **B-C)** Example calculations of CoM and Pearson’s coefficient of correlation. Elution profile for Samm50 shown for demonstration purposes. CoM is defined as the elution fraction at which the area under the curve is evenly distributed. Pearson’s coefficient of correlation is defined using standard approaches. Displayed elution profiles subject to curve smoothing as in Figure 1b-e. **D-F)** Potential changes in supercomplex composition/integrity detected by ΔCoM and Pearson’s correlation coefficient. Here, a hypothetical protein forms a supercomplex with a peak at fraction 40 in WT mice (blue). A horizontal shift in elution peak in R14^Δ/+^ mice (red) may indicate the loss of one or more subunits, affecting the apparent MW of the entire supercomplex population (D). In contrast, skewing of the peak towards lower-MW fraction (E) may indicate an altered equilibrium between a fully-assembled supercomplex and intermediate assemblies or monomers in a subpopulation. Altered Pearson’s correlation may indicate gross changes in supercomplex composition (F). **G-H**): Expected probability distributions and cutoffs used in this study. ΔCoM is expected to be normally distributed with a mean of 0 (no change, black dotted line). Probability distribution thus might be approximated with a Gaussian function, and rank-plot approximated with a cumulative Gaussian function (G). A 95^th^ percentile cutoff was chosen for this study (red dotted line). As this study is interested in disruption of high-MW protein complexes, which presumably manifest as a decrease in overall MW as illustrated in D) and E), hits with increased CoM are not considered. Probability distribution of Pearson’s correlation expected to follow an exponential decay function, and rank-plot expected to follow a log-decay function (H). A Pearson’s coefficient cutoff of ≥0.6 was chosen for this study (red dotted line).

We work under the assumption that ΔCoM will be approximately normally distributed (Figure 3g) with a mean value of 0 (i.e. no change), whereas Pearson’s coefficient is expected to follow an approximately exponential-decay probability distribution (Figure 3h). For this application, cut-offs were based on percentile (ΔCoM) and threshold value (Pearson’s). Hits can then be subjected to targeted candidate-based analysis of specific proteins of interest, or gene-ontology analysis to identify statistically-significantly overrepresented cellular components or molecular pathways (Figure 3a).

### 4. PERCOM identifies several inner-mitochondrial membrane proteins with altered elution profiles in adult 28wk-old R14^Δ/+^ mice

We applied PERCOM to identify proteins with altered distribution in R14^Δ/+^ mice. As expected, the frequency distribution of ΔCoM was centered near 0 and followed a Gaussian-like (Cauchy) distribution (Figure 4a-b) while Pearson’s coefficient followed an exponential decay frequency distribution (Figure 4c-d). A 95^th^ percentile cutoff (corresponding to ∼2x standard deviations from the mean assuming approximate Gaussian distribution) was used for ΔCoM, and a cutoff of ≥0.6 was used for Pearson’s coefficient: similar cutoffs are used in established workflows to detect complexes *de novo* by coelution^37^. We identified 296 proteins hits (out of a total of 3055 detected in both WT and R14^Δ/+^ tissue): 117 with ΔCoM, 144 with Pearson’s, and 35 with both (Figure 4e-f, Supplemental spreadsheet T3). As expected, ΔCoM proved generally suitable for identifying proteins shifted to lower-MW fractions, and Pearson’s correlation for proteins with altered elution profile shapes (Figure 4g-i).

**Figure 4:**
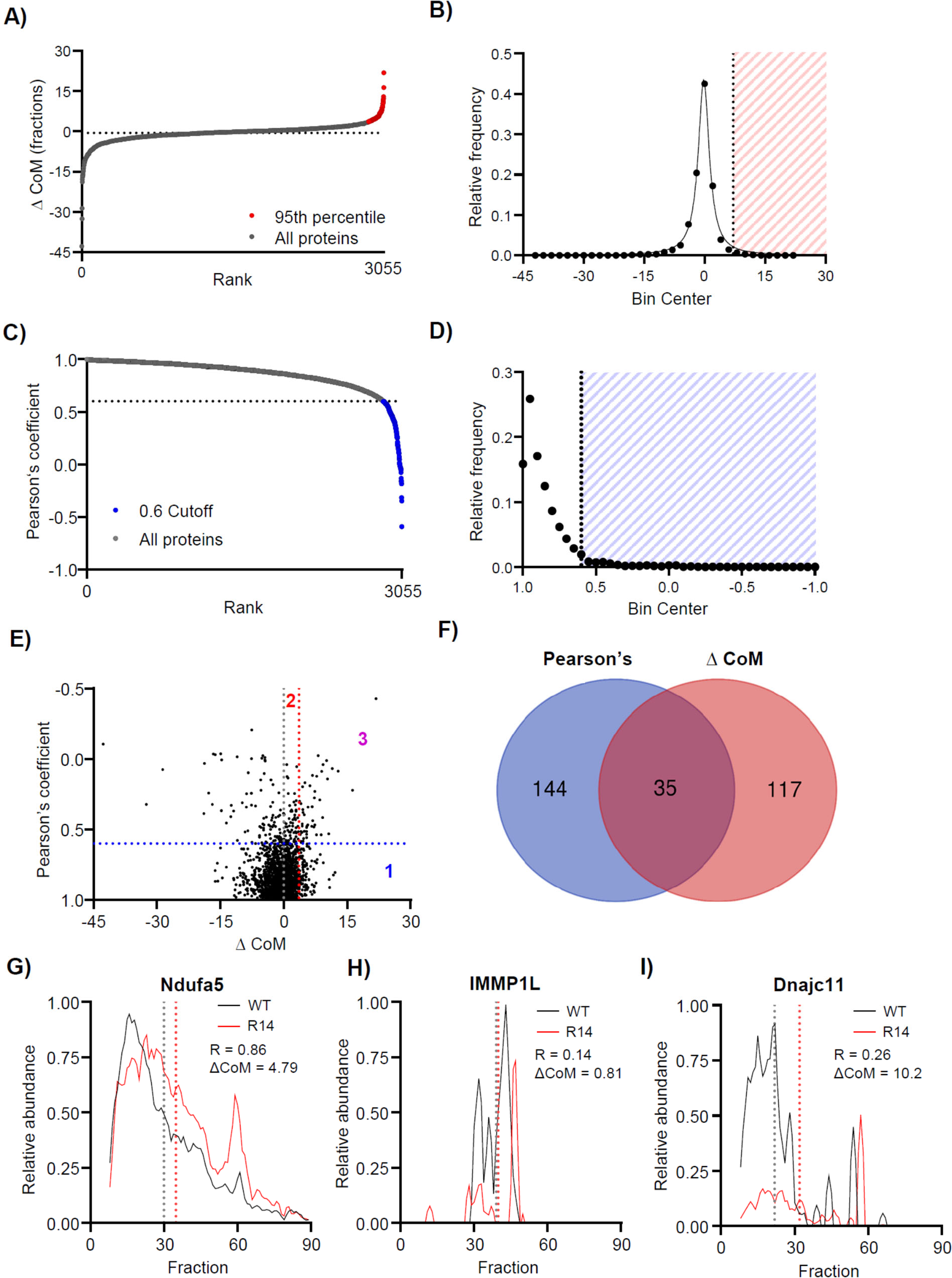
PERCOM identifies proteins with altered MW-profiles in adult 28wk-old R14^Δ/+^ mice. **A-B)** ΔCoM analysis of 28wk old WT vs. R14^Δ/+^ mouse LV tissue. Results represented as rank-plot (left) and distribution histogram (right). Distribution could be approximated by a Gaussian-like (Cauchy) distribution function (solid line). 95^th^ percentile threshold indicated in red yielded 177 hits out of 3055 proteins detected in both WT and R14^Δ/+^ tissue. **C-D)** Pearson analysis. Results represented as rank-plot (left) and distribution histogram (right). Hits (Pearson’s correlation <0.6) indicated in blue: 144 proteins out of 3055 detected. **E)** Orthogonal plot of Pearson correlation vs ΔCoM. Cutoffs indicated with dotted line. 3 quadrants are identified: hits with altered CoM (1), hits with Pearson > 0.6 (2), and hits satisfying both criteria (3). **F)** Venn diagram of hits. **G)** Example fraction profiles for a protein with altered CoM (Ndufa5), a protein with Pearson’s coefficient <0.6 (IMMP2L), and a protein meeting both criteria (Dnajc11). CoM indicated by dotted lines, and Pearson’s correlation coefficient (R) and ΔCoM shown. Experimental workflow shown in Figure 1a (N = 1). Elution profiles subject to curve smoothing as in Figure 1b-e.

### 5. Alterations in the mitochondrial respirasome supercomplexes in 28wk-old R14^Δ/+^ mice

Gene ontology analysis of PERCOM hits showed a statistically-significant enrichment of mitochondrial components (Table 1). Among affected components is the mitochondrial respirasome supercomplex (RSC), containing mitochondrial complexes CI, CIII and CIV. The coupling of mitochondrial CI, CIII and CIV proton pumping activity in RSCs allows for increased metabolic efficiency with reduced ROS production^58,59^. In cardiomyocytes, RSC assembly plays an important role in maintaining a high level of ATP synthesis and maintaining energy balance^60^. Conversely, loss of RSC integrity is associated with cardiomyopathies including DCM^61^, heart failure HF^62^ and ischemia-reperfusion injury^63^.

**Table 1:** Gene ontology overrepresentation analysis of PERCOM hits from 28wk-old WT vs R14^Δ/+^ mice (N = 1). Hits with altered CoM and Pearson’s coefficient subject to separate analysis and shown in parts 1 and 2 of the table, respectively. Mitochondrial and intercalated disk components highlighted in orange and green, respectively. Gene Ontology based on biological component performed by Panther version 18^141^. Statistical significance determined based on corrected false discovery rate (FDR) below 5% (0.05)

| 1) Change in Center of Mass (CoM) |  |  |  |
| --- | --- | --- | --- |
| GO cellular component | Fold enriched | FDR | Examples |
| MICOS complex | 92.17 | 2.14E-05 | MICOS complex subunit Mic27, Mic19, Mic10, Dnajc11 |
| SAM complex | 82.95 | 1.87E-02 | Samm50, Dnajc11 |
| HFE-transferrin receptor complex | 69.13 | 2.43E-02 | Serotransferrin, Transferrin receptor protein 1 |
| ribosomal subunit | 17.99 | 2.57E-13 | 40S ribosomal proteins S2, S3, S8, etc. 60S ribosomal proteins L24, L14, L11, etc. |
| Mitochondrial Respirasome | 12.96 | 5.11E-04 | Uqcr11, Ndufs6, Ndufa5, Uqcc3 |
| caveola | 12.70 | 5.57E-04 | Flotillin-1 and 2, Caveolin-1 and 2, Cavin-4, Nitric oxide synthase (endothelial) |
| 2) Pearson's correlation coefficient |  |  |  |
| GO cellular component | Fold enriched | FDR | Examples |
| mitochondrial inner membrane peptidase complex | > 100 | 2.00E-02 | Mitochondrial inner membrane protease subunit 1 and 2 |
| Desmosome | 29.07 | 2.26E-03 | Desmin, Desmoplakin, Desmoglein-1-alpha, Plakophilin-1 |
| Nucleosome | 19.96 | 2.07E-07 | Histone proteins (H2bc9, H4f16, H2ax, etc.) |
| cytosolic large ribosomal subunit | 19.57 | 3.36E-05 | 60S ribosomal proteins (L14, L27a, L11, etc.) |

To visualize potential RSC changes, we compared the elution profiles of the individual proteins comprising mitochondrial complexes CI-V in WT and R14^Δ/+^ mice. Constituent proteins of mitochondrial complexes CI, III and IV, but not II and V, co-eluted in a broad ≥2 MDa high-MW peak, which we interpret to correspond to the RSC (Figure 5a-e). Several CI proteins were identified as PERCOM hits and, we found the general elution profile for many CI proteins to be visibly disrupted in R14^Δ/+^ mice (Figure 5a). The profile of CIII proteins was also shifted slightly towards lower MW-fractions (Figure 5b). Complex CIV remained visually unchanged (Figure 5c). To quantify these changes, we calculated the percentage of each protein migrating above a certain MW cutoff. Cutoffs were chosen based on the apparent elution profile peaks in the WT-mouse sample: fraction #30 for RSC components (CI, CIII and CIV), and fraction #50 for CII and CIV (Figure 5a-e). We found a significant decrease in the percentage of RSC component proteins migrating above the physiological MW-fraction cutoff in R14^Δ/+^ mouse. A similar effect was noted for CV, but not CII (Figure 5f). Elution profiles for the plasma-membrane Na^+^/K^+^-ATPase (Figure 1d, Figure 5I) and the 26S proteasome (Figure 1b, Supplemental Figure S7) shown as negative controls.

**Figure 5:**
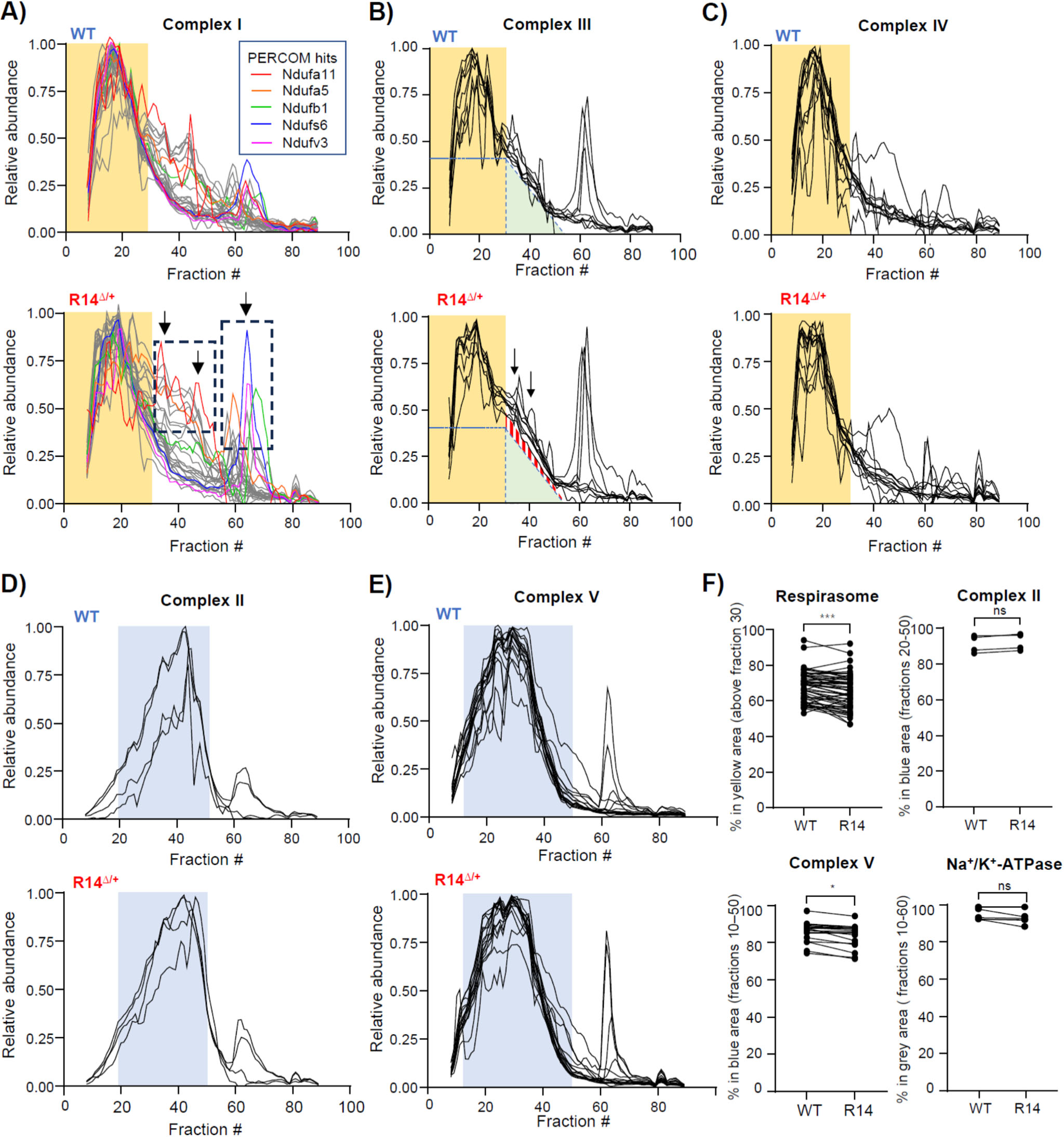
Alterations to mitochondrial respirasome supercomplexes in adult 28wk-old R14^Δ/+^ mice. **A-C)** Elution profiles of respirasome components from 28wk-old WT and R14^Δ/+^ mice. Each trace represents the elution profile of a single protein (protein IDs listed in Table 2). Experimental workflow shown in Figure 1a (N = 1). The respiratory supercomplex (RSC) contains mitochondrial complexes CI, CIII and CIV, and occupies a high-MW peak between fractions 0 and 30 (indicated with yellow underlay). **A)** Elution profiles of mitochondrial complex I proteins. Proteins identified as PERCOM hits shown in color. Areas of notable difference between WT (top) and R14^Δ/+^ mice (below) indicated with arrows and dotted boxes. **B)** Elution profiles of mitochondrial complex III proteins. In addition to the RSC peak (yellow underlay), CIII proteins also form an intermediate-MW “tail” in WT mice (green triangle). This “tail” is more pronounced in R14^Δ/+^ mice (red triangle, arrows). **C)** Elution profiles of Complex IV proteins presented as in A). **D-E)** Elution profiles of mitochondrial complexes CII and CIV not part of the RSC. Both complexes form lower-MW peaks (blue underlay) that is distinct in both apparent MW and shape from the RSC-peak shown in A-C). **F)** Quantification of apparent RSC supercomplex integrity. The area under the curve of each protein was calculated, and the percent lying within the indicated MW-fraction range plotted. Each paired data point represents the % area-under-curve of a single protein. MW-Fraction ranges correspond to the yellow and blue color underlays in panels A-E. Na^+^/K^+^ ATPase shown as a control (elution profile shown in Figure 1d). Significance determined via mixed one-way ANOVA with Šidák corrections for multiple comparison (***p < 0.001, *p < 0.05, ns = non-significant). All elution profiles subject to curve smoothing as in Figure 1b-e.

**Table 2:**
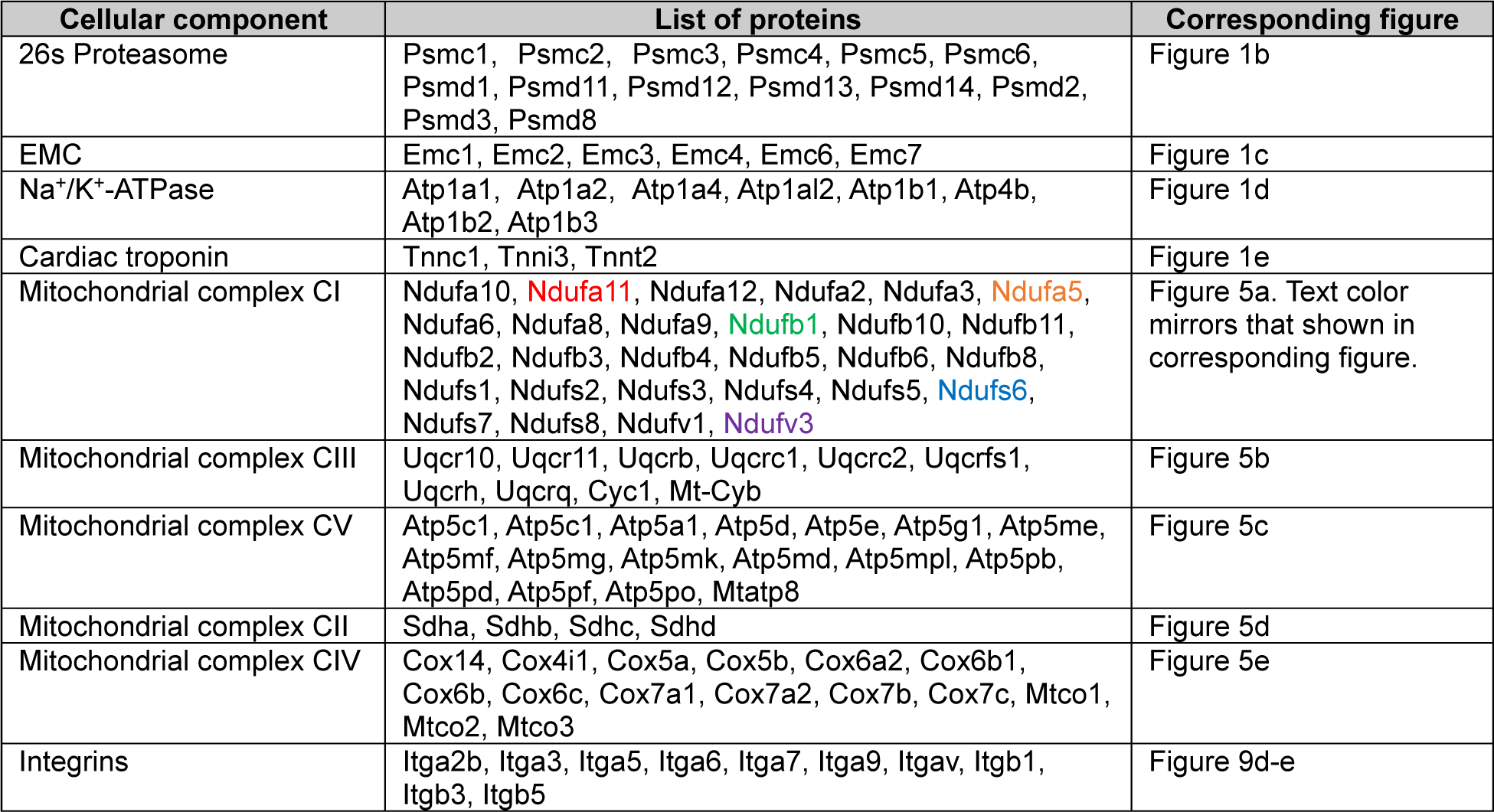
List of individual elution profiles from 28wk-old WT and R14^Δ/+^ mice shown in this work.

### 6. Reduced mitochondrial protein expression and function in 21–28-week-old R14^Δ/+^ mice

We then asked whether these changes to mitochondrial respiratory components is also reflected on the level of protein expression. LV-tissues from 21wk-week-old WT and R14^Δ/+^ mice (N = 4) were subjected to MS-based proteomics analysis using a tandem mass tag (TMT)-based isotopic labeling workflow for normalization. 3422 proteins were detected in this experiment, with 194 being significantly depleted in R14^Δ/+^ mice (Figure 6a-c). Gene ontology analysis on these depleted proteins revealed a statistically significant overrepresentation of RSC components (respiratory complex I and IV, Figure 6a, Table 3); similar statistical analysis on proteins enriched in R14^Δ/+^ mice yielded no results (Table 3). The reduction in RSC component abundances mirrors our observation of altered RSC elution profiles (Figure 5). In contrast, mitochondrial complexes CII, CIII and CV were not altered (Figure 6b).

**Figure 6:**
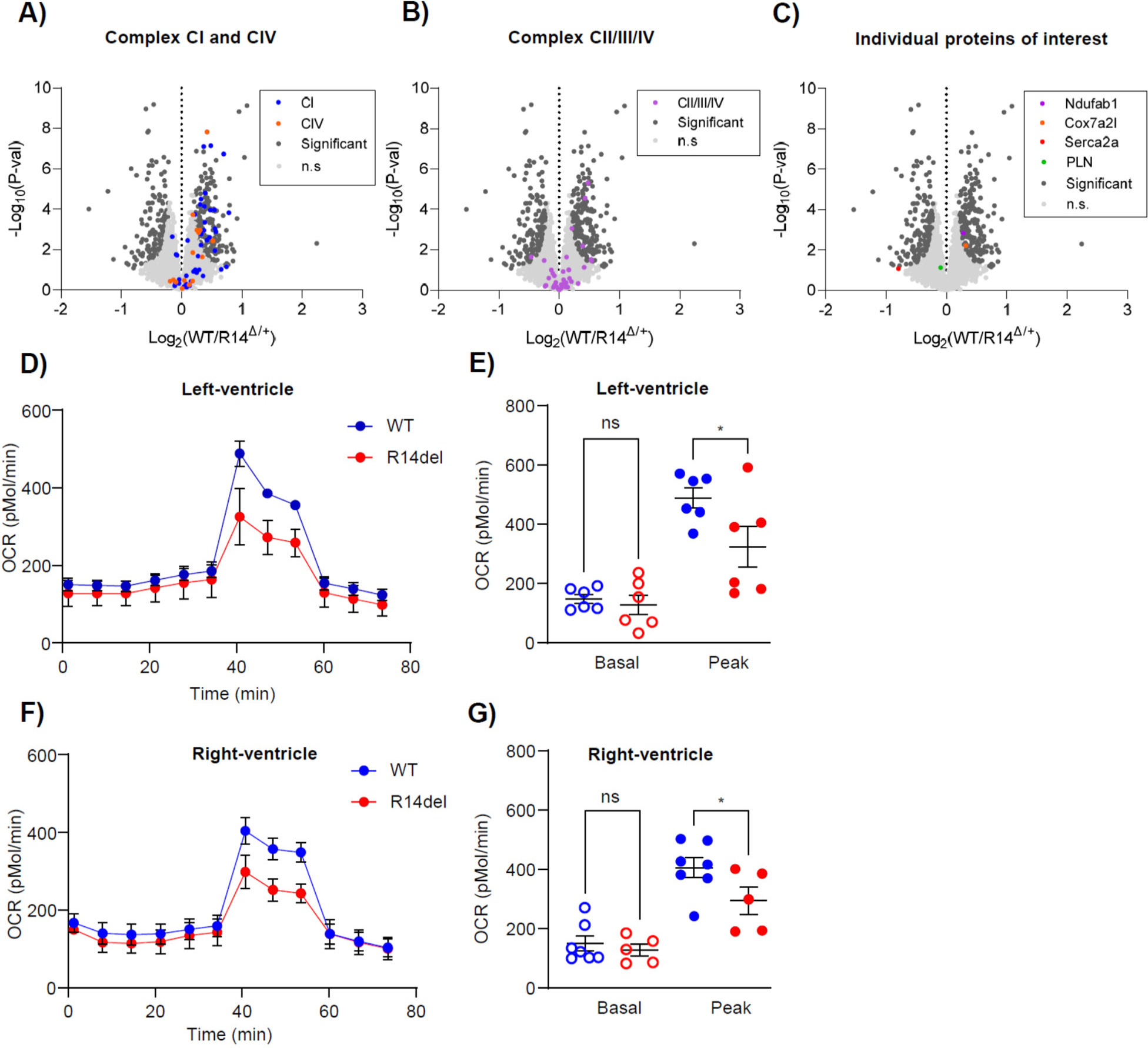
Reduced mitochondrial protein expression and function in adult 21-28wk-old R14^Δ/+^ mice. **A-C)** Global protein expression assessed in 21 wk-old ♂ WT and R14^Δ/+^ mice via bottom-up DIA-MS proteomics workflows (N = 4). Protein expression represented as a Volcano plot. Proteins with significant and non-significant (ns) changes in expression indicated with dark- and light-grey dots, respectively. **A)** Reduced expression of mitochondrial complex CI and CIV components (blue and orange, respectively). CI and CIV components were found to be significantly overrepresented amongst depleted proteins (Table 3). **B)** Expression of mitochondrial complex CII, CIII and CV components (purple). **C)** Expression of individual proteins of interest: Ndufab1 (purple), Cox7a2l (orange), SERCA2a (red) and PLN (green). **D-G)** Peak metabolic rate reduced in R14^Δ/+^ mice. Left- and right-ventricular cardiomyocytes isolated from 28wk-old WT and R14^Δ/+^ mice and Basal and peak O_2_ consumption measured by Seahorse (N = 5–7). P-values determined by one-way ANOVA with Šidák correction for multiple comparison. * p < 0.05, ns = non-significant.

**Table 3:** Gene ontology analysis of proteins with reduced expression in 21wk-old R14^Δ/+^ mouse hearts as measured by quantitative proteomics (DIA-MS, N = 4). Mitochondrial components highlighted in orange. Gene Ontology performed by Panther v. 18^141^. Statistical significance determined based on corrected false discovery rate (FDR) below 5% (0.05)

| Proteins depleted in R14<sup>Δ/+</sup> mice |  |  |  |
| --- | --- | --- | --- |
| GO cellular component | Fold enriched | FDR | Examples |
| SAM complex | 17.70 | 9.01E-03 | Samm50, Mtx2, Dnajc11 |
| mitochondrial respiratory chain complex I | 8.62 | 6.25E-09 | Ndufa1, Ndufa3, Ndufb3, Ndufs6, Ndufc2, Mtnd4, Ndufab1 |
| mitochondrial respiratory chain complex IV | 6.64 | 4.11E-02 | Cox7b, mt-Co3, Cox5b, Cox7a2l, Cox7a1, Cox4i1 |
| peroxisome | 2.88 | 5.07E-03 | Vimentin, Mul1, Pex19 |
| Proteins enriched in R14<sup>Δ/+</sup> mice |  |  |  |
| GO cellular component | Fold enriched | FDR | Examples |
| No statistically enriched components detected |  |  |  |

To-date, two proteins have been identified to play a critical role in RSC assembly: NDUFAB1^61,64,65^ and COX7A2L^66,67^. Loss of either has been shown to reduce mitochondria efficiency and increase ROS production^61,68^. Interestingly, expression of both of these assembly factors was significantly reduced in R14^Δ/+^ mice (Figure 6c), which may underlie the observed changes in RSC composition. As an interesting side-note, we observed an increase in SERCA2a expression in R14^Δ/+^ mice (Figure 6c), in contrast to recent reports showing no change in SERCA2a expression in human R14Δ-PLN patients^69^.

We next explored whether the observed RSC disruption is associated with impaired mitochondrial function. Mitochondrial function was evaluated using the Seahorse assay. While basal oxygen consumption rates (OCRs) were not significantly decreased, maximal OCR was reduced in left- and right-ventricular cardiomyocytes isolated from 28wk-old R14^Δ/+^ mice (Figure 6d-g), consistent with recent reports ^14^.

### 7. Alterations to mitochondrial respiratory supercomplexes occurs in juvenile 9wk old R14^Δ/+^ mice

To determine whether mitochondrial alterations occur at even younger ages, we performed complexome profiling on juvenile male 9wk-old WT and R14^Δ/+^ mice (N = 1, Figure 7a, Supplemental spreadsheets T4-5). As this was a confirmatory, rather than an exploratory experiment, we reduced the number of collected fractions from 89 to 47. The elution profiles of respiratory complexes CI – CV were compared (Figure 7b-g). As was the case with 28wk-old mice, we found similar alterations in the profiles for CI and CIII. CII, CIV, CV, and unrelated protein complexes such as the 26S proteasome and Na^+^/K^+^-ATPase were unchanged (Figure 7e-g, Supplemental Figure S8). These results suggest that functional and protein-level alterations at the mitochondria is an extremely early, potentially pathological change occurring in juvenile (9wk-old) animals. We observed similar changes in ventricular cardiomyocytes isolated from 12wk-old R14^Δ/+^ mice (data not shown).

**Figure 7:**
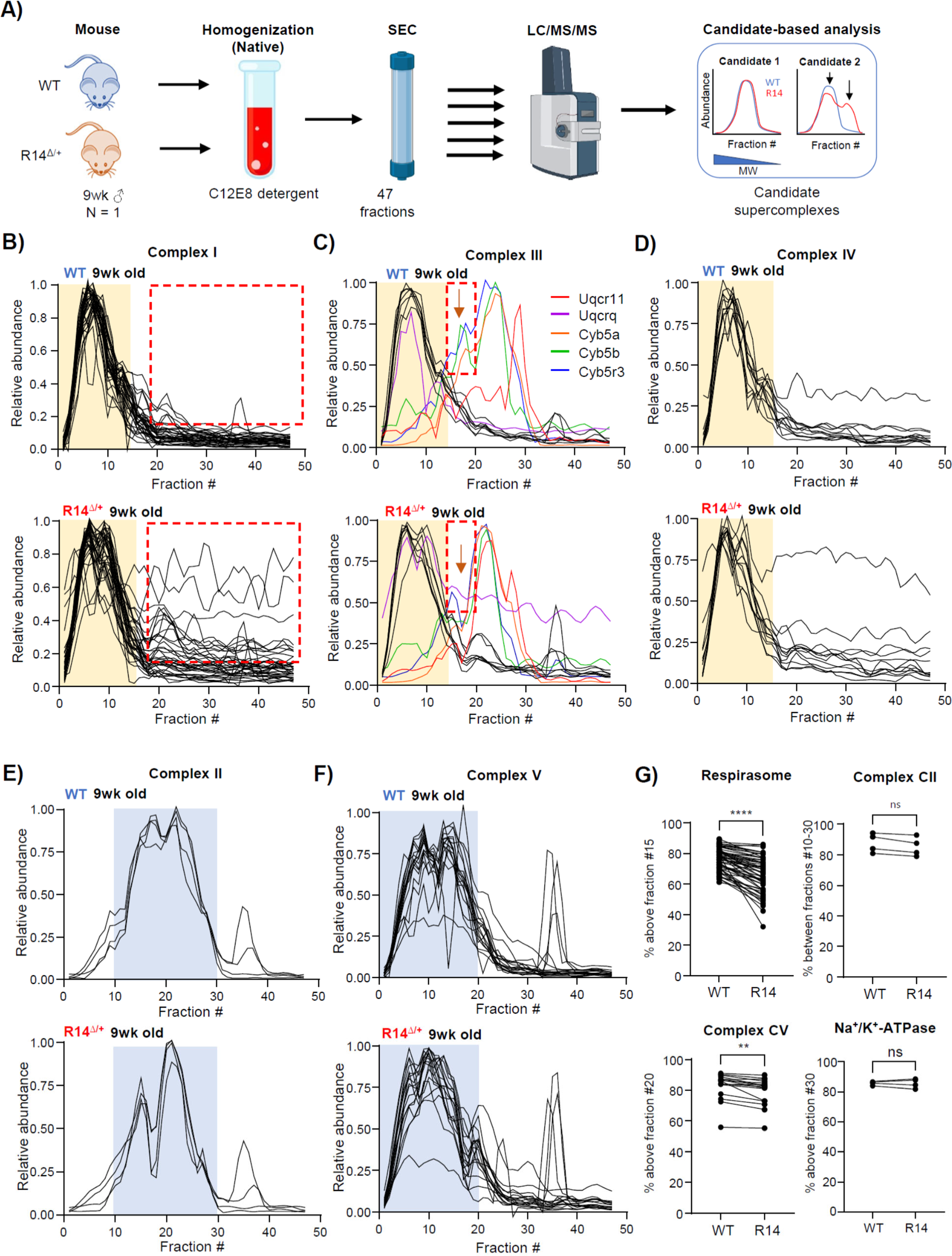
Alterations to mitochondrial respirasome supercomplexes in juvenile 9wk-old R14^Δ/+^ mice. **A)** SEC-MS workflow for 9wk-old mice. LV tissues was dissected from sibling-matched 9 wk-old ♂ WT and R14^Δ/+^ mice (N = 1). Membrane fractions were enriched by differential centrifugation, solubilized in C12E8 (0.5 mg/mg protein) and separated into 47 fractions by SEC. Each fraction was then subject to MS-based proteomics analysis. Fraction profiles of candidate proteins/protein-complexes were compared. **B-D)** Elution profiles of respirasome components from 9 wk-old WT and R14^Δ/+^ mice. Each trace represents the elution profile of a single protein. Proteins with altered elution profiles indicated by various colors. Full list of protein IDs in Table 4. Respiratory supercomplex (RSC) containing CI, CIII and CIV occupies a high-MW peak between fractions 0 and 15 (indicated with yellow underlay). Areas of notable differences highlighted by red dashed box. **E-F)** Elution profiles of mitochondrial complexes CII and CIV not part of the RSC. Both complexes form intermediate-MW peaks (blue) distinct in both shape and apparent MW from the RSC-peak shown in B-D). **G)** Quantification of apparent RSC supercomplex integrity as in Figure 5f. Elution profile for Na^+^/K^+^-ATPase shown in supplemental figure S8. All elution profiles subject to curve smoothing as in Figure 1b-e. Significance determined via mixed one-way ANOVA with Šidák corrections for multiple comparison. ****p < 0.0001, ** p < 0.01, ns = no significant difference.

**Table 4:**
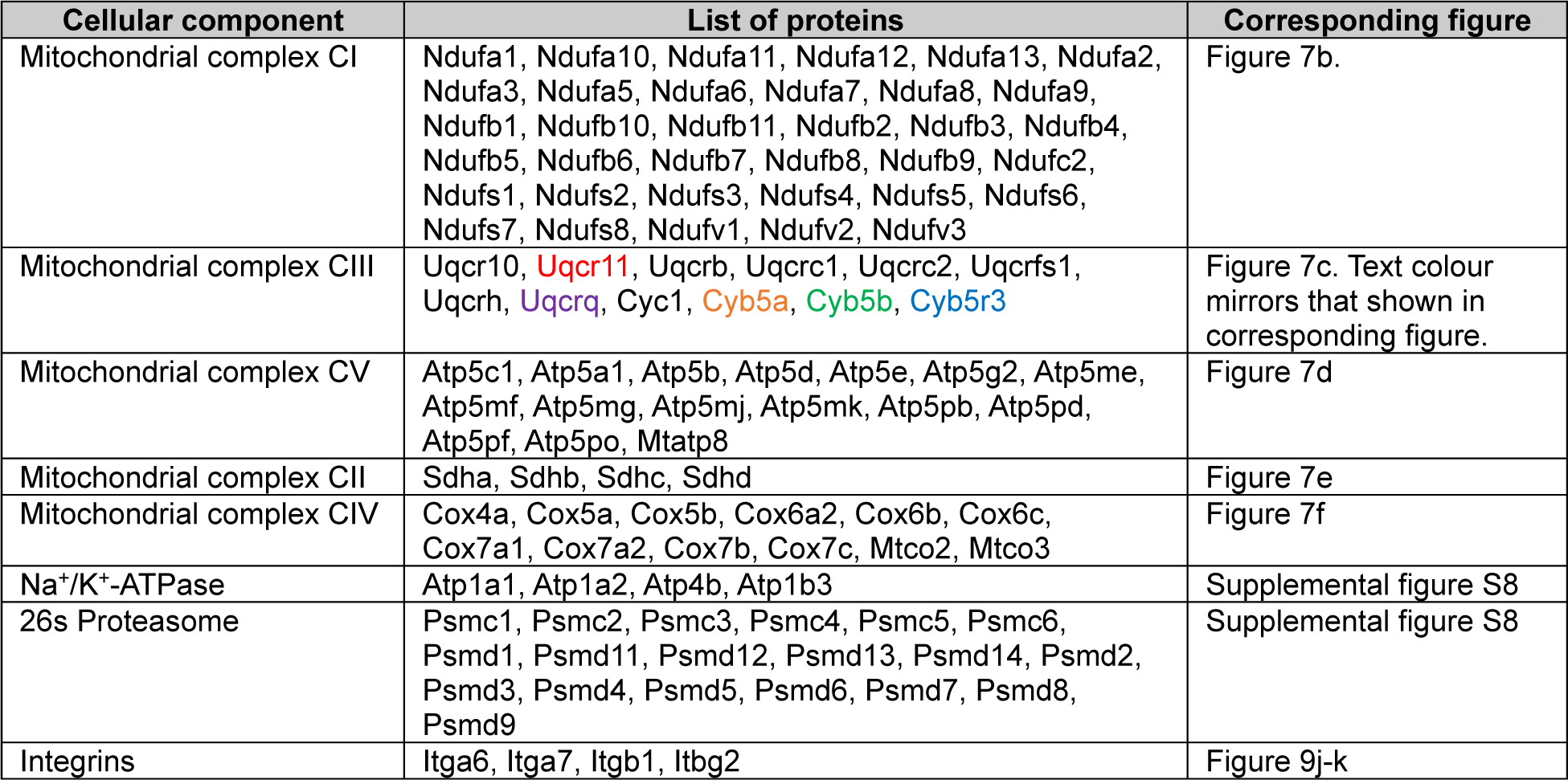
List of individual elution profiles from 9wk-old WT and R14^Δ/+^ mice shown in this work.

### 8. A mitochondrial MICOS/TOM/SAM supercomplex involved in inner-mitochondrial membrane organization is disrupted in 9wk and 28wk old R14^Δ/+^ mice

In addition to the RSC, PERCOM analysis in 28wk-old mice identified several other components involved in mitochondrial organization and protein import/sorting (Table 1). Of particular interest is the mitochondrial contact site and cristae organizing system (MICOS) complex. MICOS is an evolutionarily-conserved structure that plays a key role in mitochondrial organization. It is located primarily at cristae junctions (CJs) of the inner mitochondrial membrane (IMM)^70^ where it plays a critical role in CJ formation by providing membrane curvature^71^. It also acts as a hub for protein import and sorting through physical interactions with the translocase of the outer mitochondrial membrane (TOM) complex responsible for protein import through the OMM^72,73^ and the sorting and assembly machinery (SAM) complex responsible for membrane insertion of OMM proteins^74^. Lastly, it plays a key role the organization of respiratory complexes into the RSC^75,76^ (and may thus be functionally associated with our observed changes in the RSC).

To-date, two MICOS subcomplexes have been identified in yeast: a “core component” containing Mic60 oligomers and accessory subunit Mic19^73^, and a “Mic27 subcomplex” containing Mic10, Mic12, Mic26 and Mic27^77^. To visualize potential changes in MICOS complex assembly and/or composition, we compared the elution profiles of these components in 28wk-old mice (Figure 8a). We found that the Mic27 subcomplex was visibly altered, with 3 out of 4 detected components identified as PERCOM hits (Mic10, Mic12, Mic27, but not Mic26). In addition, Mic19, but not the core Mic60 protein was also affected. Interestingly, assembly of the Mic27 subcomplex is dependent on ER-mitochondrial contact sites, whereas self-assembly of Mic60 oligomers occurs independently of both Mic19 and ER–mitochondria contact^78^. Previous work has shown that the R14Δ-PLN negatively impacts ER-mitochondrial contact sites in cardiomyocytes differentiated from R14^Δ/+^ hiPSC-CMs^50^. Whether ER-mitochondrial contact sites are disrupted in adult transgenic mice remains unknown; however, this would be consistent with our finding of altered Mic27 subcomplex assembly.

**Figure 8:**
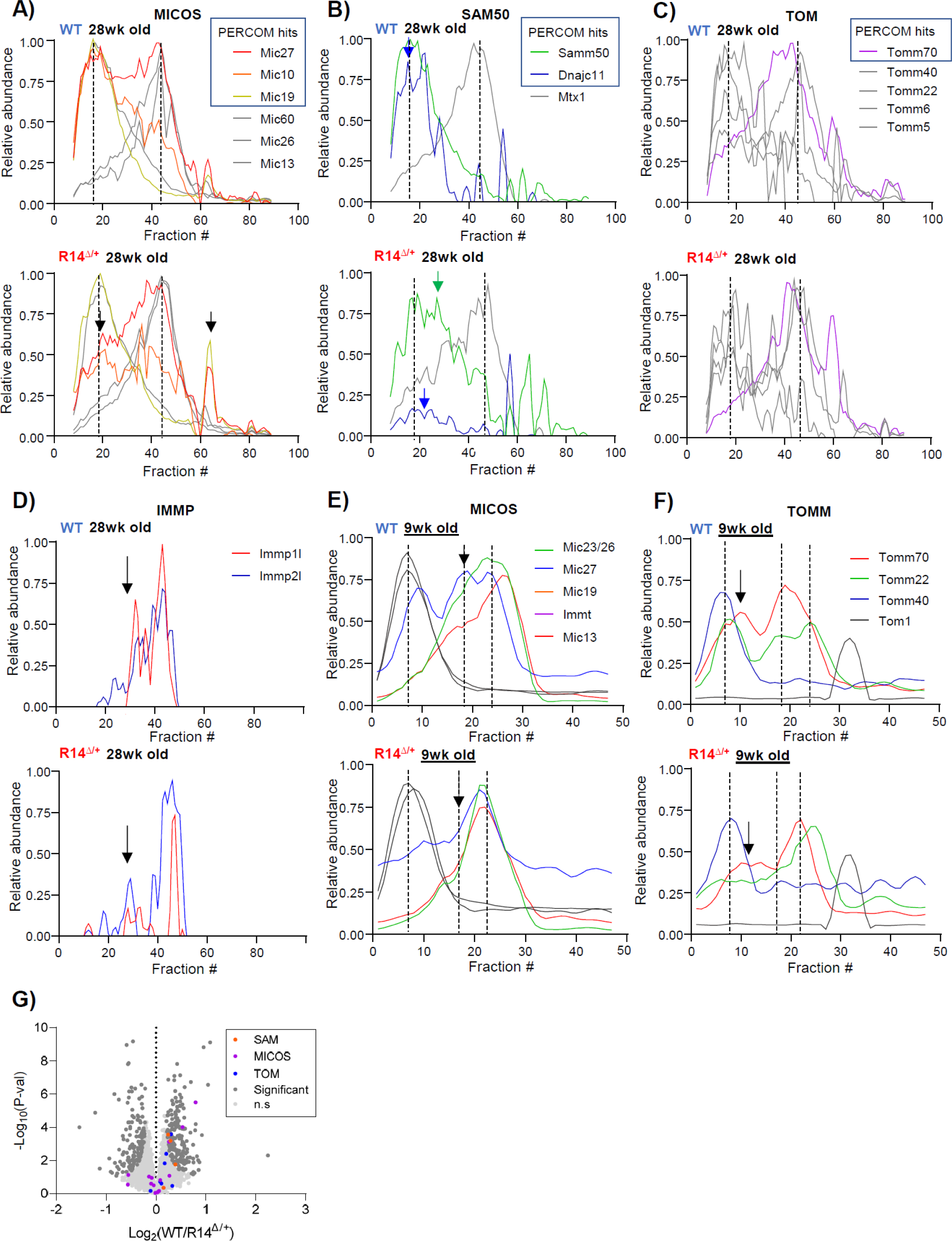
Alterations to protein complexes involved in mitochondrial organization in28wk adult and 9wk juvenile R14^Δ/+^-mice. **A-C)** A putative MICOS/SAM/TOM supercomplex is disrupted in adult 28wk-old R14^Δ/+^ mice. Elution profiles of proteins comprising the MICOS, SAM50 and TOM complexes shown. All components at-least partially co-elute as two high-MW peaks (dotted vertical lines), suggesting that MICOS, SAM50 and TOM associate as a supercomplex. PERCOM hits indicated with colored lines; unaltered proteins in gray. Areas of notable difference indicated by arrows. **D**) Alterations to the inner mitochondrial membrane peptidase (IMMP) complex in adult 28 wk-old R14^Δ/+^ mice. The IMMP complex consists of two proteins (Immp1l and Immp2l), both of which were identified as PERCOM hits (red and blue traces, respectively). Areas of notable difference indicated by arrows. Immp1l trace shown previously in Figure 4h. **E-F)** Alterations to MICOS and TOM complexes in juvenile 9 wk-old R14^Δ/+^ mice. Co-eluting peaks indicated with dotted lines as in panels A-C. SAM50 complex components were not detected in this experiment. Areas of notable difference indicated by arrows. **G)** Decreased expression of proteins involved in inner-mitochondrial membrane (IMM) organization. Data and plot shown in Figure 6a-c re-colored to highlight MICOS, Sam50 and TOM protein complex components. Subcellular compartment annotation retrieved from UniprotKB. Experimental workflows shown in Figure 1a and 7a (N = 1). All elution profiles subject to curve smoothing as in Figure 1b-e.

The MICOS complex co-ordinates with other import/organizational machinery such as the TOM^72,73^ and SAM ^74^ complexes. MICOS, SAM and TOM components at-least partially co-eluted in both WT and R14^Δ/+^ mice (Figure 8a-c), consistent with the formation of a higher-order MICOS/SAM/TOM supercomplex^79^. Notably, components of the SAM complex were identified as being statistically enriched among our PERCOM hits (Table 1), raising the possibility that R14Δ-PLN may alter not only the MICOS complex, but also this putative MICOS/SAM/TOM supercomplex. Indeed, the elution profiles of both SAM and TOM components were altered in 28wk-old R14Δ mice (Figure 8b-c). Lastly, we examined the inner mitochondrial membrane peptidase (IMMP) complex. This assembly is responsible for proteolytic maturation of proteins targeted to the intermembrane space^80^. The IMMP complex was identified as a PERCOM hit (Table 1), and both IMMP components (IMMP1 and IMMP2) displayed radically altered elution profiles in 28wk-old R14Δ mice (Figure 8d).

We then asked whether changes in the putative MICOS/SAM/TOM supercomplex and IMMP complex occur in juvenile 9wk-old R14^Δ/+^ mice. While we were unable to detect components of the SAM and IMMP complexes in our more limited dataset, we did find changes to the elution profiles of MICOS and TOM components (Figure 8e-f). Lastly, the quantitative MS-based proteomics analysis shown earlier also revealed SAM complex components to be significantly depleted in 21-wk-old R14^Δ/+^ mice (Table 3, Figure 8g). Taken together, our findings suggest that R14Δ-PLN impairs assembly/stability of key machinery involved in mitochondrial protein import and organization.

### 9. Elution profiles of intercalated disk proteins are altered in 9wk and 28wk old R14^Δ/+^ mice

In addition to mitochondrial supercomplexes, gene ontology analysis of PERCOM hits identified desmosomes as another potentially impacted structure (Table 1). Desmosomes are an important component of the cardiac intercalated disk (ICD), which mediates mechanical and electrical coupling between adjacent cardiomyocytes, fibroblasts and macrophages^81–85^. Within ICDs, mechanical coupling is mediated by the aforementioned desmosomes containing desmosomal cadherins such as Desmoglein-2 (Dsg2), Desmoplakin (Dsp) and plakoglobin (Jup), alongside adherens junctions composed of classical Cadherin (Cdh2) and Catenin-α/β^81^. In direct vicinity, electrical coupling is mediated by gap junctions composed of connexion hemichannels, of which connexin-43 (Cx43) is the major ventricular isoform^81^. Mutations to proteins at the desmosome^86–88^, adherens junction^89–92^ or gap junction^93,94^ are associated with conduction slowing, ACM and DCM.

The impact of R14Δ-PLN on cardiac desmosomes was evaluated by comparing elution profiles between WT and R14^Δ/+^ adult (28wk-old) mice (Figure 9). Numerous desmosomal proteins were identified by PERCOM (Tables 1); however, only a subset (Dsg2, Jup) was detected as high-MW complexes (Figure 9a). Nonetheless, both of these proteins were shifted towards low-MW fractions in R14^Δ/+^ mice (Figure 9a, f). We then evaluated the integrity of the adherens and gap junction complexes, where we found a similar shift towards lower-MW fractions (Figure 9b-c, f). In contrast, the MW profiles of α/β Integrin, another family of cell-surface adhesion proteins ^95^, was unchanged (Figure 9d-f). Lastly, we evaluated the integrity of intercalated disk complexes in juvenile (9wk-old) mice. We were unable to detect desmosomal proteins in high-MW fractions (data not shown); however, all detected adherens proteins Cadherin-2 (Cdh2) and Catenin-α1 (Ctnna1), as well as Cx43 showed altered elution profiles (Figure 9g-i). Integrins were once again unaffected (Figure 9j-k).

**Figure 9:**
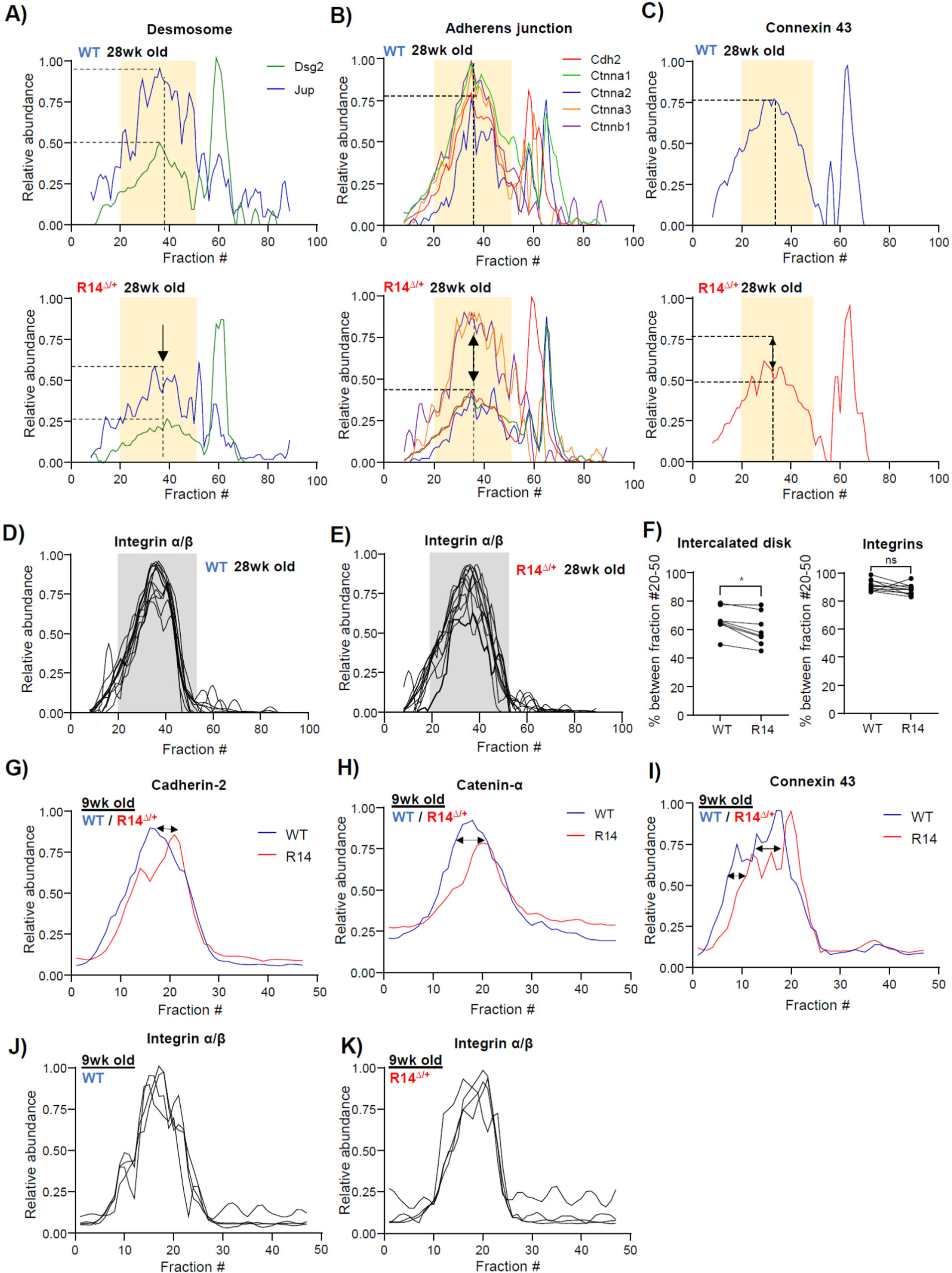
Alterations to intercalated-disk components in 28wk-old adult and 9wk-old juvenile R14^Δ/+^-mice. **A-C)** Elution profiles for structural proteins of the desmosome (A), adherens junction (B) and gap junction (C). All components coelute as a high-MW peak centered on fraction 38 (vertical dashed line). The distribution of a subset of proteins within this peak is reduced in R14^Δ/+^-mice, as indicated by dashed horizontal lines and arrows. **D-E)** Elution profiles of Integrin α/β components. Full list of protein IDs in Table 2. **F)** Fraction of detected ICD and Integrin proteins eluting within MW-fraction 20-50, indicated with yellow and gray underlay, respectively. Data represented as in Figure 5f. Significance determined via mixed one-way ANOVA with Šidák corrections for multiple comparison (*p < 0.05, ns = no significant difference). **G-I**) Elution profiles of all detected adherens proteins (Cadherin-2, Catenin-A) and gap junction protein connexin-43 from juvenile 9 wk-old WT and R14^Δ/+^ mice. Elution profiles of all proteins shifted towards lower-MW fractions in R14^Δ/+^-animals (arrows). **J-K)** Elution profiles of Integrin α/β are not altered in 9 wk-old R14^Δ/+^ mice. Full list of protein IDs in Table 4. Experimental workflows shown in Figure 1a and 7a (N = 1).

Overall, these results suggest that intercalated disk architecture may be altered at an early stage of R14Δ-PLN cardiomyopathy. This may contribute to the observed electrophysiological abnormalities such as a reduced conduction velocity observed in R14^Δ/+^ mouse hearts^7,39^ and increased arrhythmia risk in both transgenic mice^7,39,96^ and human patients^3^.

## Discussion

Here, we describe the development of improved data acquisition and analysis workflows for the profiling of high-MW assemblies in cardiac tissues. Using these workflows, we identified alterations in the elution profiles of several key mitochondrial and ICD components. These findings could be observed in presymptomatic R14^Δ/+^ mice as young as 9 weeks of age. Cardiomyocytes have a uniquely high demand for ATP due to their constant pumping activity, and are dependent on mechanical and electrical connectivity between adjacent cells to facilitate force transduction and propagation of electrical excitation; thus, our observed alterations of mitochondrial and ICD supercomplexes could have particularly severe consequences to cardiac function. While previous studies have shown impaired mitochondrial function in hiPSC-CM models^50^, our findings are among the first to make this observation in a presymptomatic animal system^14^, and the first to demonstrate alterations at the molecular and protein levels.

### Molecular-level mitochondrial alterations are a hallmark of early-stage R14Δ-PLN cardiomyopathy

Here, we present three lines of evidence demonstrating mitochondrial alterations as a hallmark of early-stage R14Δ-PLN cardiomyopathy in presymptomatic (9-28 wk-old) R14^Δ/+^ mice: altered elution profiles of the RSC and a putative MICOS/SAM/TOM supercomplex (via complexome profiling), reduced expression of RSC and SAM component proteins (via MS-based proteomics analysis) and reduced maximal OCR (via Seahorse). Previous work has shown reduced mitochondrial function in patient-derived hiPSC-CM models^50^ and, only recently, in presymptomatic R14^Δ/+^ mice^14^. Thus, our results are not only among the first reports of mitochondrial alterations as an early-disease event in presymptomatic animals (consistent with it being a causative mechanism contributing to R14Δ-PLN pathogenesis), but also provide the first direct evidence of changes at the molecular/protein level. Observed changes to mitochondrial protein elution profiles were relatively subtle (Figures 5, 7 and 8), suggesting minor shifts towards smaller sub-assemblies rather than widespread supercomplex disruption. This is consistent with recent work showing no significant change to mitochondrial membrane potential and cytosolic reactive oxygen species (ROS) content in 8-12 wk-old R14^Δ/+^ mice^14^ and the overall lack of significant cardiomyopathy in these animals^7,14^, and together paint a picture of subtle changes to mitochondrial organization and function.

How R14Δ-PLN (an integral SER-membrane protein) can exert influence in the mitochondria remains an open question. R14Δ-PLN is associated with various Ca^2+^-handling defects, including a hyperdynamic phenotype^14^ and increased diastolic [Ca^2+^]^97^. Altered Ca^2+^ dynamics may lead to mitochondrial Ca^2+^-overload resulting in damage to mitochondrial supercomplexes^98^ (Figure 10) which is a key determinant of heart failure pathology^99^. The recent observation that acute reduction of cytosolic or SER [Ca^2+^] results in a worsening rather than an improvement in mitochondrial function ^14^, as well as failure to detect increased ROS generation in R14^Δ/+^ mouse ventricular myocytes, a hallmark of Ca^2+^-overload^98^, at rest would appear to argue against this. Alternatively, impaired Ca^2+^-reuptake may require compensation via increased Ca^2+^ secretion via the plasma membrane Ca^2+^-ATPase (PMCA) and/or the Na^+^/Ca^2+^-exchanger (NCX, with increased Na^+^ influx being countered by increased Na^+^/K^+^-ATPase activity)^100^, resulting in increased ATP demand and metabolic imbalance. In addition, increased UPR activity^13^ and protein aggregation^7,12^ may also increase ATP demand (ex. via the activity of ATP dependent chaperones and proteasomal activities)^101,102^ (Figure 10).

**Figure 10:**
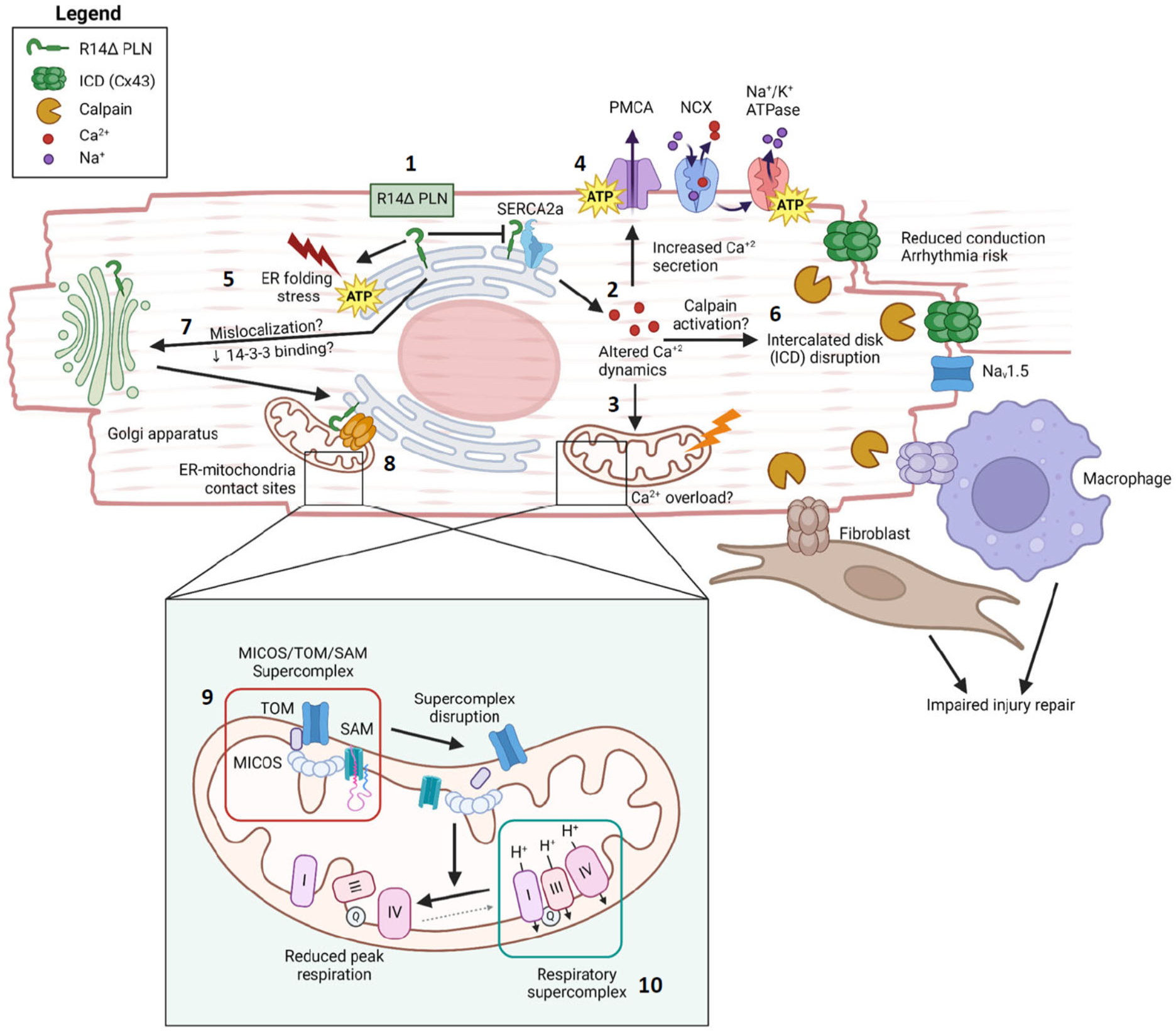
Potential cellular model of observed molecular changes. At the SER, R14Δ-PLN is associated with SERCA2a superinhibition (1), resulting in abnormal Ca^2+^-dynamics (2). This may directly lead to mitochondrial Ca^2+^-overload (3). Increased ATP-dependent Ca^2+^-secretion via plasma-membrane transporters may also increase ATP demand, leading to further mitochondrial stress (4). ER-folding stress, UPR activation, and upregulation of ATP-dependent proteostasis pathways may also contribute to increased energy demand (5). Lastly, intercalated disks (ICDs) mediate critical contacts with neighboring cardiomyocytes, macrophages and fibroblasts, and are vital for conduction propagation and injury repair, respectively. ICD components (Connexin-43, Cx43 illustrated as example) may be degraded by aberrant Calpain activation (6). Loss of diarginine motif may result in ER-exit (due to loss of the ER-retention signal and/or binding of phospho-adaptor protein 14-3-3) and trafficking to post-ER compartments (7). Mislocalized R14Δ-PLN may aberrantly associate with ER-mitochondrial sites, resulting in further mitochondrial effects (8). Inset: at the mitochondria, we describe a putative MICOS/TOM/SAM supercomplex (9) that is involved in import/insertion of mitochondrial proteins, organization of mitochondrial membrane topology, and stabilization of respiratory supercomplexes containing complexes CI/CIII/CIV. Alterations to this supercomplex in R14^Δ/+^ PLN hearts lead to a partial disruption of the respiratory supercomplex and reduced respiratory efficiency (10).

We also speculate that mislocalization of a subpopulation of R14Δ-PLN to post-ER compartments, previously observed in both HEK293 cells and transgenic mouse models^103,104^, may contribute to mitochondrial dysfunction. The R14Δ mutation disrupts a R13/R14 diarginine motif that may act as an ER-retention signal^105,106^ and 14-3-3 binding site^107^; this may result in the mislocalization of a subpopulation (small enough to evade detection by microscopy, Figure 2) to peripheral cellular compartments^105,108^. Interestingly, our complexome profiling data shows that PLN co-elutes with a subset of ER-mitochondrial tethering proteins, including Calnexin, BCAP31 and VAPA/B (Supplemental figure S9), suggesting some degree of crosstalk (Figure 10). In addition, the mitochondrial transmembrane protein TMEM 126B was identified as a hit in our PERCOM analysis (Supplemental spreadsheet T3). TMEM 126B plays a key role in the assembly of mitochondrial complex CI via the Mitochondrial Complex I Assembly (MCIA) complex^28^. TMEM126B showed increased tendency to comigrate with PLN in 28wk-old R14^Δ/+^ mice compared to WT controls (Supplemental Figure S9); it was not detected in our 9wk-old cohort dataset. As TMEM126B may span both the inner and outer mitochondrial membranes, this provides a potential direct physical link between PLN and IMM supercomplex integrity.

### Intercalated disks may represent a new site of R14Δ-PLN action

Previous reports observed a decrease in conduction velocity in 12–16wk-old R14^Δ/+^ mice hearts^7,39^, which may contribute to the R14Δ-ACM phenotype. Due to the complete absence of fibrosis at this age^7,11^, these conduction abnormalities have been initially attributed to altered Ca^2+^-homeostasis^7^. ICDs mediate electromechanical coupling between adjacent cardiomyocytes^109^, macrophages^82^ and fibroblasts^84,85^ . In addition to acting as physical portals, intercalated disks also recruit voltage-gated cation channels such as Na_v_1.5 to propagate depolarization at the gap junction to the rest of the cell^110,111^. Mutations to intercalated disk proteins such as desmoglein-2, Plakophillin-2 (PKP2) and plakoglobin are associated with decreased conduction velocity and ACM^112,113^. In addition, contact between cardiomyocytes, macrophages and fibroblasts is involved in response to cardiac injury^83,84^, and ICD disruption has also been recently implicated in DCM pathogenesis^114^. Here, we provide preliminary evidence that intercalated disk supercomplexes are altered in R14^Δ/+^ mice, which may contribute to the ACM/DCM phenotype (Figure 10). Further functional and (ultra)structural experiments should be done to establish this as a potential mechanism of R14Δ-PLN pathogenesis.

A potential mechanism linking R14Δ-PLN to ICD alterations is aberrant activation of Calpain, a peripheral Ca^2+^-dependent protease implicated in ICD disruption following myocardial infraction^115,116^. It is possible that altered Ca^2+^-homeostasis in R14Δ-PLN hearts may also result in Calpain overactivation and ICD degradation (Figure 10). The potential for aberrant degradation of other Calpain targets in R14Δ-PLN hearts, such as the SER-PM tethering protein Junctophilin-2^117–119^ also deserves future investigation.

### Mitochondrial and intercalated disk supercomplexes represent potential therapeutic targets for early intervention

A critical component of our work is the observation of disrupted ICD and mitochondrial supercomplexes in juvenile (9wk-old) R14^Δ/+^ mice, significantly prior to the onset of cardiomyopathy at 18 months of age^7,11^. Targeting these pathways at a very young age may represent an effective therapeutic strategy that can be delivered prior to the onset of clinical symptoms. Recent work has shown AAV-mediated overexpression of mitochondrial complex CI core subunit S1 (Ndufs1) and assembly factor Ndufab1 to be cardioprotective against ischemia-reperfusion injury in mice^61,120^. These gene therapy approaches may be of use in treating R14Δ-CM. In particular, Ndufab1 overexpression enhances assembly of the RSC^61^, which is a specific alteration seen in R14^Δ/+^ mice. Alternatively, several mitochondrial-targeting drugs such as MitoQ (a mitochondria-targeted antioxidant)^121,122^ and Elamipretide (a small molecule inhibitor of ROS formation)^123^ have been trialed for a wide range of diseases including Alzheimer’s^124^ and Parkinson’s disease^124^. While preclinical and clinical testing has yielded mixed results^125–129^, the potential application to R14Δ-PLN remains unexplored. In particular, the very early onset of mitochondrial alterations may make preventative treatment particularly effective (conversely, the inability to treat prior to onset of clinical symptoms may have limited the apparent effectiveness of these compounds in prior clinical trials)^127^. Therapeutic approaches targeting the ICD are less established. Should increased and/or aberrant Calpain activity be detected in R14^Δ/+^-PLN hearts (potentially leading to degradation of ICD components, Figure 10), testing of Calpain inhibitors may be warranted.

### Application of SEC-DIA-MS based complexome profiling workflows in cardiac samples

Previous complexome profiling studies on cardiac tissue employed digitonin-based solubilization of membrane and/or mitochondrial proteins followed by BNE-based fractionation and MS-based proteomics measurement^22,40^. We made several critical improvements to this data acquisition workflow: **1)** *Choice of detergent*. C12E8 was found to be superior to digitonin for the solubilization of cardiac membrane protein complexes (Supplemental Figures S1-S2). In addition, C12E8 is a synthetic detergent that is commercially available at high purity and homogeneity, whereas digitonin represents a complex mix of glycosides isolated from *Digitalis purpurea*, resulting in both lower purity and increased lot-to-lot variability^130^. **2)** *High MW range SEC-based fractionation*. This study is, to our knowledge, the first application of SEC-based fractionation to cardiac samples. Compared to BNE-PAGE, SEC offers potentially-enhanced resolution at very high (here, up to 7.5 MDa) MW ranges and enhanced reproducibility^43^. **3)** *Quantitative DIA-MS analysis using an empirical spectral library.* Previous work relied upon *in-silico* tryptic digests to generate peptide libraries for DIA data extraction. Here, we generated a dedicated spectral library specific for adult C57BL/6J mouse ventricular tissue in order to improve bottom-up proteomic depth of analysis^131^. More generally, DIA-MS has been demonstrated to increase both the speed and the reproducibility of CP experiments. Taken together, our SEC-DIA-MS workflow offers significant improvements for complexome profiling analysis in cardiac samples. This is illustrated by our ability to detect not only routinely-studied mitochondrial complexes and supercomplexes, but also cardiac-specific structures such as the SER Ca^2+^-handling supercomplex (Figure 2a), voltage-gated Ca^2+^ channel and K^+^-channel oligomers (Supplemental figure S1c), and intercalated disk components (Figure 9).

### PERCOM: an accessible and unbiased protein-centric data analysis workflow able to detect subtle changes in specialized tissue-specific supercomplexes

In addition to the improved data acquisition strategy, we also developed PERCOM as a novel data analysis workflow. PERCOM bypasses several issues associated with using curated protein complex databases for ground-truth (underrepresentation of tissue- or cell-type-specific structures, underrepresentation of higher-order supercomplexes with fluid composition, incomplete coverage of the human proteome) and can, furthermore, be implemented using basic spreadsheet software, making it broadly accessible to the scientific public.

It should also be noted that PERCOM was able to identify proteins with subtly-altered elution profiles in a presymptomatic heterozygous animal model^7,11^. This highlights the sensitivity of our workflow, and demonstrates its suitability for use in disease models with no/mild phenotypes where only subtle alterations might be expected. There is a growing appreciation that many critical functions in excitable cell types, such as Ca^+2^-signalling, excitation propagation and membrane contacts, occur with nanoscale resolution via specialized supercomplexes or membrane nanodomains^22,132^. We believe that our SEC-DIA-MS CP workflow coupled with PERCOM represents an important tools for studying higher-order protein complexes underlying these nanoscale functions in specialized cell-types, and identifying alterations in mild/asymptotic disease models.

### Conclusions and study limitations

Here, we present an optimized workflow for SEC-MS complexome profiling of mouse ventricular tissue. We developed a novel data analysis workflow (PERCOM) to ameliorate the underrepresentation of cardiac-specific entries in available protein complex databases. SEC-MS based CP workflows coupled with PERCOM identified several mitochondrial and intercalated disk supercomplex components with altered elution profiles in presymptomatic 9-wk and 28-wk old R14^Δ/+^mice. Mitochondrial changes were associated with reduced expression of mitochondrial oxidative phosphorylation proteins and maximal OCR, as evaluated using MS-based proteomics and Seahorse, respectively.

This study presents two CP datasets: one from 28wk-old male mice (Figures 1-6, 8 and 9), and one from 9wk-old male mice (Figures 7-9). The experimental cohort for each experiment was a single R14^Δ/+^ animal and a sibling-matched WT-control (N = 1, Figures 1 and 7). Due to the large number of MS-based proteomics measurements required per animal (47 or 89 fractions per sample, each measured in technical duplicate), biological replicate measurements were cost- and time-prohibitive. The same cost issue also precluded experiments on female animals. Nonetheless, our core findings were observed in both datasets despite the differences in both age and cohort, indicating biological reproducibility of our results. Lastly, while our observations at the mitochondria are supported by multiple lines of evidence (complexome profiling, global proteomics, Seahorse), as well as a body of previous work in both hiPSC-CMs^50^ and transgenic mice models^14^, our observations at the intercalated disk are admittedly limited to a single readout with no prior art. We believe further research into this topic, including orthogonal functional and (ultra)structural validation, to be an important direction for future work.

## Methods and Materials

### Animal Studies

Generation and characterization of the R14Δ-PLN knockin mouse line was described previously^7^. All experimental cohorts consist of sibling-matched male WT and R14^Δ/+^ mice unless otherwise noted. Animal experiments are performed in accordance to directive 2010/63/EU of the European Parliament and in keeping with NIH guidelines. Use of R14Δ-PLN mice is covered by animal protocol 21/3698, and additional WT-animals for assay development by institutional animal protocols 11/2 and 22/19. All protocols were reviewed and approved by the institutional animal committee of the University Medical Center Göttingen and approved by the veterinarian state authority (LAVES, Oldenburg, Germany). Experimental animals are housed in the Universitätsmedizin Göttingen Zentrale Tierexperimentelle Einrichtung (ZTE) and colonies are maintained at the Max Planck Institute for Multidisciplinary Sciences (Göttingen).

### Dissection of ventricular tissue and isolation of enriched membrane fractions

Male WT and R14^Δ/+^ mice were anaesthetized with isoflurane and sacrificed by cervical dislocation. Whole hearts were dissected, cannulated at the aorta and profused for 4 minutes with a modified Ca^+2^-free Krebs buffer via Langendorf setup at a flow rate of 4ml/min ^133^. Flash-frozen left- and right-ventricular tissues were thawed, minced and resuspended in sucrose homogenization buffer (250mM sucrose, 20mM imidazole, 6mM EDTA, 6mM Tris HCl pH 6.8) supplemented with EDTA-free protease inhibitor cocktail (Roche). Tissues were disrupted using potter homogenizer (2x 25 strokes at 2000rpm), followed by passage through a 27G needle (10x). Homogenates were centrifuged at 1,000g (10 minutes, 4°C) to remove debris, and enriched membrane fractions were pelleted by centrifugation at 100,000g (1h, 4°C). Membrane fractions were solubilized for at-least 30 minutes at 4°C in sucrose homogenization buffer containing either digitonin (6mg/mg protein), Triton X-100 (0.01mg/mg protein), NP-40 (0.5mg/mg protein) or C12E8 (0.5mg/mg protein).

### Generation of empirical spectra library and DDA/DIA-MS proteomics analysis

50 µg of enriched membrane fractions were solubilized in either SDS (2%) or C12E8 (0.5 mg/mg protein) and purified by a brief SDS-PAGE run (Nu-PAGE 4-12%, Invitrogen). Lanes were excised, reduced, alkylated, and digested with trypsin as previously described^134^. Tryptic digestion was quenched using 1% trifluoracetic acid and peptides were separated into 12 fractions by basic reverse-phase liquid chromatography (äkta pure, Bruker). Liquid chromatography (LC)-coupled tandem MS/MS analysis was performed using a hybrid timed ion mobility-time of light mass spectrometer (timsTOF Pro 2, Bruker) coupled to a nanoflow chromatography system (Ultimate nanoSRLC, Thermo Fisher Scientific) equipped with a reversed phase-C18 column (Aurora Elite 250×0.075 mm, IonOpticks) and employing a linear 2-37% acetonitrile versus 0.1% formic acid gradient at a flow rate of 250 nl/min. Samples were analyzed on in either DDA mode (one technical replicate) or DIA mode (two technical replicates). DDA analysis was performed in Parallel Accumulation−Serial Fragmentation (PASEF) mode^135^ with 10 scans per topN acquisition cycle. Multiple charged precursors were selected based on their position in the *m/z*–ion mobility plane and isolated at a resolution of 2 Th for *m/z* ≤ 700 and to 3 Th for *m/z* > 700 with a target MS/MS ratio of 20,000 arbitrary units. Dynamic exclusion was set to 0.4 min. DIA analysis was performed in diaPASEF mode^136,137^using a customized 8×2 window acquisition method from *m/z* from 100 to 1,700 and from 1/K_0_ from 0.7 - 1.5 to include the 2^+^/3^+^/4^+^ population in the *m/z*–ion mobility plane. The collision energy was ramped linearly as a function of the mobility from 59 eV at 1/K_0_ = 1.5 Vscm^−2^ to 20 eV at 1/K_0_ = 0.7 Vscm^−2^. The spectral library was built with Spectronaut software version 16.3 (Biognosys) by matching data against the Uniprot KB mouse reference proteome (01/2021).

### BNE-based complexome profiling for detergent selection

Membrane fractions from seven 28w-old ♂ WT mice were pooled and solubilized in the indicated detergent. 35-50 µg of solubilized membrane fractions were loaded onto pre-cast Blue Native-PAGE gels (3-12% Bis-Tris minigels, Invitrogen) and run using standard protocols^138^. Lanes were cut into 35 equidistant fractions using a custom stainless-steel cutter. All fractions were reduced, alkylated, digested with trypsin and dried by SpeedVac (Thermo) in 96 well microplates as previously described^134^. Fractions were spiked with iRT standard peptides (1/100 diluted, Biognosys) and subject to LC-MS/MS analysis in technical duplicates on a hybrid quadrupole-orbitrap mass spectrometer (Q Exactive, Thermo Fisher Scientific) hyphenated to a nanoflow chromatography system (Easy-nLC 1000, Thermo Fisher Scientific) equipped with a homemade analytical column (C-18aq, 3µm, 200×0.075mm, Dr. Maisch) and precolumn (Reprosil C18aq, 5µm, 20×0.15 mm, Dr. Maisch). Separation was achieved using a 5% to 35% acetonitrile gradient in 0.1% formic acid. Samples were measured in technical duplicate in DIA mode. Briefly, MS1 spectra were collected in the range of 350-1250 *m/z* at 70,000 resolution (FWHM), a maximum injection time of 50ms and a target AGC of 1 x 10^6^. Subsequently, 11 variable size DIA windows were analyzed using the following settings: default charge 3^+^, resolution 35,000 (FWHM), maximum injection time 115 ms, AGC target 3x 10^6^, fixed first mass 200 *m/z*. Precursor fragmentation was achieved using a stepped Normalized Collision Energy regime of 26/28/30%. Protein concentration in each fraction was measured by UV absorbance at 280nm prior to analysis (Supplemental figure S1). Spectral data were analyzed using Spectronaut software version 16.3.

### SEC-based complexome profiling in adult 28w-old mice

Membrane fractions were isolated from 28 wk-old ♂ WT and R14^Δ/+^ mice (N = 1) as described above. Approximately 120 µg of solubilized membrane fractions were fractionated into 89 fractions using an äkta pure chromatography system (Cytiva) equipped with a Bio-SEC-5 1000Å analytical column (300×7.8 mm, Agilent) and a SEC-5 1000 Å guard column (50×4.6 mm, Agilent). Commercial gel filtration standards (Bio-Rad) were employed for molecular weight calibration of the SEC runs. SEC was performed on ice with an isocratic flow rate of 0.5 ml/min for 36.8 minutes using a buffer containing 5 mM Tris-HCl, 1 mM EDTA, 150 mM KCl, 0.005 mg/ml C12E8 solution (pH 7) ^44^. Following fractionation, additional Tris-HCL pH 8.8 (20-25 mM) and SDC (0.2% w/v) was added and reduction, alkylation and tryptic digestion using standard protocols^32^. Fractions were spiked with iRT standard peptides (Biognosys) as described above and separated using a linear 12.5 min gradient of 4-32% aqueous acetonitrile versus 0.1% formic acid on a nanoflow chromatography system (Ultimate nanoRSLC, Thermo Fisher Scientific) using a reversed phase-C18 column (PepSep Fifteen, 150×0.150 mm, Bruker) at a flow rate of 850 nl min^-1^. Peptides were measured in technical duplicates using a hybrid timed ion mobility-time of flight mass spectrometer (timsTOF Pro 2, Bruker). MS/MS analysis was done in diaPASEF mode as described above. Protein concentration in each fraction was measured by UV absorbance at 280 nm. The first 8 fractions (#1 to 8) did not contain sufficient protein concentration to justify measurement and were thus discarded from further analysis (Supplemental Figure S3). Spectral data were analyzed using Spectronaut software version 16.3.

### SEC-based complexome profiling in juvenile 9w-old mice

Membrane fractions were isolated from 9w-old ♂ WT and R14^Δ/+^ mice (n=1) as described above. Approximately 120 µg of solubilized membrane fractions were fractionation into 47 fractions by SEC as described above. Fractions were spiked with iRT standard peptides and analyzed in technical duplicates via LC-MS/MS on a hybrid quadrupole-orbitrap mass spectrometer (Q Exactive, Thermo Fisher Scientific) hyphenated to a nanoflow chromatography systems (Easy-nLC 1000 UHPLC, Thermo Fisher Scientific) as described above. Protein concentration in each fraction was measured by UV absorbance at 280nm (Supplemental Figure S8). Analysis of spectral data was performed using Spectronaut software version 16.3 (Biognosys).

### Refinement of complexome profiling data and generation of protein abundance matrices

Following spectral analysis, data was arranged into protein abundance matrices, with rows representing individual protein IDs, columns representing fraction number, and cell values containing the detected abundance (Figure 3). A Pearson’s correlation cutoff of ≥0.6 between technical duplicates was applied to identify and eliminate proteins with poor technical reproducibility, with an exception being made for RyR2 due to its biological importance and a prime candidate for interrogation. In this case, fractions where the techical replicate measurements of protein abundance differed by more than 25% of maximum were discarded from further analysis. Protein abundance values (expressed as peptide counts) were converted into relative abundances (expressed as fraction of the maximal abundance across the fractions).

### Quantification of protein complexes defined by the CORUM protein complex database

Detection and quantification of protein complexes defined within the CORUM 4.0 protein complex database was performed using a custom R-software package (mCP). In brief: protein detection was implemented via detecting co-elution of complex components (coelution being measured by Pearson’s correlation coefficient). A ≥0.81 Pearson’s coefficient cutoff was determined using Monte-Carlo simulations (185 simulations) with a target false-discovery rate of 5%. At the time of writing, a manuscript describing this R-script package is in preparation (preprints available upon reasonable request).

### Calculation of ΔCoM and Pearson’s correlation coefficient

The elution fraction representing the center of distribution (CoM) was calculated using the following equation:

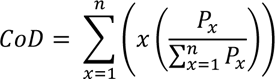

Where x represents elution fraction number, n represents the total number of fractions, and P represents the relative abundance of the protein in that fraction.

Pearson’s coefficient of correlation was calculated using the following equation^139^:

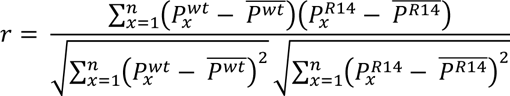

Where P^WT^ and P^R^^14^ represent the relative abundance of the protein in the indicated elution fraction number in WT and R14^Δ/+^ samples, respectively.

### MS-based proteomics analysis

Left-ventricular cardiomyocytes were isolated from 21w-old ♂ WT and R14^Δ/+^ mice (n=4) as described previously^133^. Briefly, whole hearts from 28 wk-old ♂ WT and R14^Δ/+^ mice were cannulated and perfused with modified Krebs buffer containing 2 mg/ml collagenase (Worthington) for 10 min. Digested heart tissues were dissected using a scalpel and disrupted by gentle pipetting. Myocytes were isolated by differential sedimentation. Left- and right-ventricles were dissected from digested hearts and disrupted by pipetting. Cardiomyocytes were homogenized by pipetting for 5 minutes at 4°C in lysis buffer (0.05% v/v NP40, 50 mM HEPES, 150 mM NaCl, 50 mM NaF, 200 μM Na_3_VO_4_, pH 7.5) supplemented with 1 mM PMSF and protease inhibitor cocktail (Roche). Lysates were then reduced and alkylated with 10 mM DTT and 100 mM of 2-Chloroacetamide (30 min each step at 30°C) and digested with trypsin (1/40 w/w ratio) at 37°C for 14 h. TMT labelling (1 µg reagent per µg peptide, 50 µg peptide per reaction) was performed at 25°C for 30 minutes and quenched with 5% hydroxylamine (15min at 25°C). Desalting and peptide concentration was performed using reversed phase Sep-Pak C18 cartridges. Labelled peptides were fractionated into 12 fractions using basic pH reversed phase liquid chromatography on a reversed phase-C18 column (Hypersil Gold, 150×2.1 mm, Thermo Fisher Scientific) using a 0% - 90% acetonitrile gradient. LC-MS/MS was performed in technical duplicate using an tribrid mass spectrometer (Orbitrap Fusion, Thermo Fisher Scientific) hyphenated to a nanoflow chromatography system (Ultimate nanoRSLC, Thermo Fisher Scientific) equipped with a reversed phase-C18 chromatography column (Reprosil C18aq, 1.9 μm, 280×0.075 mm, Dr. Maisch).

Spectral data were analyzed using MaxQuant software version 1.6 (Max Planck Institute of Biochemistry). Datasets were analyzed against the UniProtKB mouse reference proteome (06/2019, 55485 entries). Under group specific parameters, the TMT labeling search was made with MS3 reporter ion with the 6plex TMT labeling on lysine and N-terminal residue. Reporter mass tolerance was by default set to 0.003 Da and a maximum of two missed cleavages allowed. Variable modifications include protein N-terminal acetylation and methionine oxidation and fixed modifications contain cysteine carbamidomethylation. Peptides and proteins were reduced using a 1% peptide spectrum match (PSM) false discovery rate (FDR) and a 1% protein FDR determined by the decoy database. Only the razor/unique peptides were used for peptide identification and quantitative calculations.

Protein group data were processed by Perseus v1.6.15.0 (Max-Planck-Institute of Biochemistry). Site only, reverse, and contaminant peptides were removed from the dataset and missing values were imputed using a normal distribution (width: 0.3, downshift: 1.8). Invalid values were then excluded. Empty columns were removed. Rows were categorically annotated as wild type and mutant respectively for each reported intensity corrected and the data was transformed with log2(x). For the removal of the batch effect within TMT 6plex, data normalization was done in Perseus using width adjustment quartile in program normalization.

### Super-resolution confocal microscopy

Cardiomyocytes were isolated from mouse left-ventricle as described above and seeded onto fibronectin-coated glass cover slips. PLN, SERCA2a and RyR2 were labelled using the following antibodies: monoclonal mouse α-PLN (Thermo), polyclonal rabbit α-SERCA2a (Badrilla) and rabbit α-RyR2 (Sigma). Secondary antibodies conjugated to STED-compatible fluorophores (STAR 580 or STAR 635P, Abberoir) were used for labelling. Images were acquired using a Leica TCS SP8 STED system using an HC PL APO C2S 100x/1.40 oil immersion lens. Image analysis was performed on FiJi using a custom macro similar to those used in our earlier works^22,40^.

### Mitochondrial functional assays

Left- and right-ventricular cardiomyocytes were isolated as described above and seeded onto fibronectin-coated Seahorse XF cell culture plates (5000 cells/well). Measurement of oxygen consumption rate (OCR) was done in a Seahorse XF96e Extracellular Flux Analyzer (Agilent Technologies) according to the instructions provided by the manufacturer. The measurement was done in XF-DMEM buffer supplemented with 1 mM pyruvate, 2 mM glutamine, and 10 mM glucose. Basal oxygen consumption rate was measured as well as respiration following the addition of 3 µM Oligomycin, 1.5 µM CCCP, and 0.5 µM Antimycin/Rotenone to determine oxygen consumption under differing metabolic conditions.

### Gene Ontology, statistical analysis, and presentation of data

Functional and subcellular-component classifications of proteins were extracted from Uniprot^140^. Enrichment and overrepresentation analysis was performed by Panther 18.0^141–143^. Statistical analysis was performed using GraphPad Prism 9 (Dotmatics) or Microsoft Excel (Microsoft). Statistical significance was determined using two-tailed T-tests (with Bonferroni correction for multiple comparisons), one-way or two-way ANOVA (with Šidák corrections) as indicated in the respective figure legends. Figures 1a, 7a and 10, as well as supplemental figure S1a generated using art assets from Biorender (biorender.com) with appropriate permissions and licenses (available upon reasonable request).

### Patents

There are no granted or applied for patents relating to this manuscript.

## Supporting information

Supplemental spreadsheet 1: CP data for 28wk old WT mouse

Supplemental spreadsheet 2: CP data for 28wk old R14del/+ mouse

Supplemental spreadsheet 3: PERCOM hits (28wk old mice)

Supplemental spreadsheet 4: CP data for 9wk old WT mouse

Supplemental spreadsheet 2: CP data for 9wk old R14del/+ mouse

## Acknowledgements

The authors would like to acknowledge Dana El-Chami, Rebecca Rosen Falk, Tobias Kohl, Brigitte Korff, Lisa Neuenroth, Timo Schulte, Birgit Schumann and Fabio Trebini for technical support; Dr. Daniel Kownatzki-Danger for preliminary work and setup of this project, and Christiane Schulz for administrative support. Mass spectrometry measurements were supported by the University Medical Center Göttingen (UMG) Core Facility Proteomics.

## Author Contributions

Conceptualization: B.F., C.L. and S.E.L. Generation of R14Δ-mouse line: H.H.W.S. and R.A.DB. Isolation of mouse material: B.F and H.A. Native homogenization method development: H.A. SEC method development: H.A. Complexome profiling measurements (SEC): H.A. and C.L. Complexome profiling measurements (BNE): H.A. Mass-spectrometry based quantitative proteomics: S.K. and C.L. Immunofluorescence microscopy: S.J. Measurement of oxygen consumption rate: A.B., T.G. and P.R. Overall data analysis: B.F. and H.A. Drafting of manuscript: B.F. Revision of manuscript: B.F., H.A., H.H.W.S., R.A. DB., H.U., C.L. and S.E.L, All authors have read and agreed to the published version of the manuscript.

## Data availability

Custom computer script for quantification of protein complexes using external databases is currently being prepared for publication (Amedei *et al.*). The mass spectrometry proteomics data have been deposited to the ProteomeXchange Consortium via the PRIDE^144^ partner repository with the dataset identifier PXD050383.

## References

1. Fabiato, A. & Fabiato, F. Calcium Release from the Sarcoplasmic Reticulum. Circ Res 40, 119–129 (1977).

2. James, P., Lnui, M., Tada, M., Chiesit, M. & Carafoli, E. Nature and site of phospholamban regulation of the Ca 2 + pump of sarcoplasmic reticulum. Nature 342, 90–92 (1989).

3. Schmitt, J. P. et al. Dilated Cardiomyopathy and Heart Failure Caused by a Mutation in Phospholamban. Science (1979) 299, 1410–1413 (2003).

4. Haghighi, K. et al. Human phospholamban null results in lethal dilated cardiomyopathy revealing a critical difference between mouse and human. Journal of Clinical Investigation 111, 869–876 (2003).

5. Haghighi, K. et al. A mutation in the human phospholamban gene, deleting arginine 14, results in lethal, hereditary cardiomyopathy. Proc Natl Acad Sci U S A 103, 1388–1393 (2006).

6. Hof, I. E. et al. Prevalence and cardiac phenotype of patients with a phospholamban mutation. Netherlands Heart Journal 27, 64–69 (2019).

7. Eijgenraam, T. R. et al. The phospholamban p.(Arg14del) pathogenic variant leads to cardiomyopathy with heart failure and is unreponsive to standard heart failure therapy. Sci Rep 10, (2020).

8. Verstraelen, T. E. et al. Prediction of ventricular arrhythmia in phospholamban p.Arg14delmutation carriers–reaching the frontiers of individual risk prediction. Eur Heart J 42, 2842–2850 (2021).

9. Vafiadaki, E., Haghighi, K., Arvanitis, D. A., Kranias, E. G. & Sanoudou, D. Aberrant PLN-R14del Protein Interactions Intensify SERCA2a Inhibition, Driving Impaired Ca2+ Handling and Arrhythmogenesis. Int J Mol Sci 23, (2022).

10. Qin, J. et al. Structures of PKA–phospholamban complexes reveal a mechanism of familial dilated cardiomyopathy. Elife 11, (2022).

11. Stege, N. M. et al. DWORF Extends Life Span in a PLN-R14del Cardiomyopathy Mouse Model by Reducing Abnormal Sarcoplasmic Reticulum Clusters. Circ Res 133, 1006–1021 (2023).

12. Eijgenraam, T. R. et al. Protein Aggregation Is an Early Manifestation of Phospholamban p.(Arg14del)-Related Cardiomyopathy: Development of PLN-R14del-Related Cardiomyopathy. Circ Heart Fail 14, e008532 (2021).

13. Feyen, D. A. M. et al. Unfolded Protein Response as a Compensatory Mechanism and Potential Therapeutic Target in PLN R14del Cardiomyopathy. Circulation 144, 382–392 (2021).

14. Maniezzi, C. et al. Early consequences of the phospholamban mutation PLN-R14del+/− in a transgenic mouse model. Acta Physiologica (2024) doi:10.1111/apha.14082.

15. Nooren, I. M. A. & Thornton, J. M. Diversity of protein-protein interactions. EMBO Journal 22, 3486–3492 (2003).

16. Marsh, J. A. & Teichmann, S. A. Structure, dynamics, assembly, and evolution of protein complexes. Annu Rev Biochem 84, 551–575 (2015).

17. Vercellino, I. & Sazanov, L. A. The assembly, regulation and function of the mitochondrial respiratory chain. Nat Rev Mol Cell Biol 23, 141–161 (2022).

18. Seflova, J. et al. Fluorescence lifetime imaging microscopy reveals sodium pump dimers in live cells. Journal of Biological Chemistry 298, (2022).

19. Blackwell, D. J., Zak, T. J. & Robia, S. L. Cardiac Calcium ATPase Dimerization Measured by Cross-Linking and Fluorescence Energy Transfer. Biophys J 111, 1192–1202 (2016).

20. Roderick MacKinnon. Determination of the subunit stoichiometry of a voltage activated potassium channel. Nature 350, 323–325 (1991).

21. Frank, R. A. & Grant, S. G. Supramolecular organization of NMDA receptors and the postsynaptic density. Current Opinion in Neurobiology vol. 45 139–147 Preprint at 10.1016/j.conb.2017.05.019 (2017).

22. Alsina, K. M. et al. Loss of Protein Phosphatase 1 Regulatory Subunit PPP1R3A Promotes Atrial Fibrillation. Circulation 140, 681–693 (2019).

23. Wu, H. Di et al. Ultrastructural remodelling of Ca2+ signalling apparatus in failing heart cells. Cardiovasc Res 95, 430–438 (2012).

24. De Groot, J. R. & Coronel, R. Acute ischemia-induced gap junctional uncoupling and arrhythmogenesis. Cardiovasc Res 62, 323–334 (2004).

25. Calore, M., Lorenzon, A., De Bortoli, M., Poloni, G. & Rampazzo, A. Arrhythmogenic cardiomyopathy: a disease of intercalated discs. Cell and Tissue Research vol. 360 491–500 Preprint at 10.1007/s00441-014-2015-5 (2015).

26. Páleníková, P., Minczuk, M. & Rebelo-Guiomar, P. Protocol to study human mitochondrial ribosome using quantitative density gradient analysis by mass spectrometry and complexome profiling analysis. STAR Protoc 4, 102605 (2023).

27. Páleníková, P. et al. Quantitative density gradient analysis by mass spectrometry (qDGMS) and complexome profiling analysis (ComPrAn) R package for the study of macromolecular complexes. Biochim Biophys Acta Bioenerg 1862, (2021).

28. Heide, H. et al. Complexome profiling identifies TMEM126B as a component of the mitochondrial complex i assembly complex. Cell Metab 16, 538–549 (2012).

29. Olinares, P. D. B., Ponnala, L. & Van Wijk, K. J. Megadalton complexes in the chloroplast stroma of Arabidopsis thaliana characterized by size exclusion chromatography, mass spectrometry, and hierarchical clustering. Molecular and Cellular Proteomics 9, 1594–1615 (2010).

30. Kristensen, A. R., Gsponer, J. & Foster, L. J. A high-throughput approach for measuring temporal changes in the interactome. Nat Methods 9, 907–909 (2012).

31. Kirkwood, K. J., Ahmad, Y., Larance, M. & Lamond, A. I. Characterization of native protein complexes and protein isoform variation using sizefractionation-based quantitative proteomics. Molecular and Cellular Proteomics 12, 3851–3873 (2013).

32. Bludau, I. et al. Complex-centric proteome profiling by SEC-SWATH-MS for the parallel detection of hundreds of protein complexes. Nat Protoc 15, 2341–2386 (2020).

33. Heusel, M. et al. Complex-centric proteome profiling by SEC - SWATH - MS . Mol Syst Biol 15, (2019).

34. Michalak, W., Tsiamis, V., Schwämmle, V. & Rogowska-Wrzesińska, A. ComplexBrowser: A tool for identification and quantification of protein complexes in large-scale proteomics datasets. Molecular and Cellular Proteomics 18, 2324–2334 (2019).

35. Tsitsiridis, G., et al. CORUM: the comprehensive resource of mammalian protein complexes-2022. Nucleic Acids Res 51, D539–D545 (2023).

36. Giese, H., et al. NOVA: A software to analyze complexome profiling data. Bioinformatics 31, 440–441 (2015).

37. Stacey, R. G., Skinnider, M. A., Scott, N. E. & Foster, L. J. A rapid and accurate approach for prediction of interactomes from co-elution data (PrInCE). BMC Bioinformatics 18, (2017).

38. Gorka, M. et al. Protein Complex Identification and quantitative complexome by CN-PAGE. Sci Rep 9, (2019).

39. Raad, N. et al. Arrhythmia Mechanism and Dynamics in a Humanized Mouse Model of Inherited Cardiomyopathy Caused by Phospholamban R14del Mutation. Circulation 144, 441–454 (2021).

40. Peper, J. et al. Caveolin3 Stabilizes McT1-Mediated Lactate/Proton Transport in Cardiomyocytes. Circ Res 128, E102– E120 (2021).

41. Szczepanowska, K. et al. A salvage pathway maintains highly functional respiratory complex I. Nat Commun 11, (2020).

42. Hevler, J. F., et al. Molecular characterization of a complex of apoptosis-inducing factor 1 with cytochrome c oxidase of the mitochondrial respiratory chain. PNAS 118, (2021).

43. Barth, H. G., Boyes, B. E. & Jackson, C. Size Exclusion Chromatography. Anal. Chem. 68, 445–466 (1996).

44. Santos, H. L., Lamas, R. P. & Ciancaglini, P. Na,K-ATPase Solubilized Using Only C 12 E 8. Brazilian Journal of Medical and Biological Research vol. 35 (2002).

45. Le Maire, M., Lind, K. E., Jorgensen, K. E., Rgigaard, H. & Mallerg, J. V. Enzymatically Active Ca2+ATPase from Sarcoplasmic Reticulum Membranes, Solubilized by Nonionic Detergents. Journal of Biological Chemistry 263, 7051–7058 (1978).

46. Levesque, I., Juliano, B. R., Parson, K. F. & Ruotolo, B. T. A Critical Evaluation of Detergent Exchange Methodologies for Membrane Protein Native Mass Spectrometry. J Am Soc Mass Spectrom 34, 2662–2671 (2023).

47. Juliano, B. R., Keating, J. W. & Ruotolo, B. T. Infrared Photoactivation Enables Improved Native Top-Down Mass Spectrometry of Transmembrane Proteins. Anal Chem 95, 13361–13367 (2023).

48. Yoshikawa, H. et al. Efficient analysis of mammalian polysomes in cells and tissues using Ribo Mega-SEC. Elife (2018) doi:10.7554/eLife.36530.001.

49. Cleary, S. R., et al. Dilated cardiomyopathy variant R14del increases phospholamban pentamer stability, blunting dynamic regulation of cardiac calcium handling. BioRxiv (2023) doi:10.1101/2023.05.26.542463.

50. Cuello, F. et al. Impairment of the ER/mitochondria compartment in human cardiomyocytes with PLN p.Arg14del mutation. EMBO Mol Med 13, (2021).

51. Meldal, B. H. M. et al. Complex Portal 2022: New curation frontiers. Nucleic Acids Res 50, D578–D586 (2022).

52. Van Strien, J. et al. COmplexome Profiling ALignment (COPAL) reveals remodeling of mitochondrial protein complexes in Barth syndrome. Bioinformatics 35, 3083–3091 (2019).

53. Mariuzza, R. A., Agnihotri, P. & Orban, J. The structural basis of T-cell receptor (TCR) activation: An enduring enigma. Journal of Biological Chemistry vol. 295 914–925 Preprint at 10.1074/jbc.REV119.009411 (2020).

54. Lingaraju, G. M. et al. Crystal structure of the human COP9 signalosome. Nature 512, 161–165 (2014).

55. Söhl, G. & Willecke, K. Gap junctions and the connexin protein family. Cardiovasc Res 62, 228–232 (2004).

56. Insel, P. A. et al. Caveolae and lipid rafts: G protein-coupled receptor signaling microdomains in cardiac myocytes. In Annals of the New York Academy of Sciences vol. 1047 166–172 (New York Academy of Sciences, 2005).

57. Letts, J. A. & Sazanov, L. A. Clarifying the supercomplex: The higher-order organization of the mitochondrial electron transport chain. Nat Struct Mol Biol 24, 800–808 (2017).

58. Maranzana, E., Barbero, G., Falasca, A. I., Lenaz, G. & Genova, M. L. Mitochondrial Respiratory Supercomplex Association Limits Production of Reactive Oxygen Species from Complex I. Antioxid Redox Signal 19, (2013).

59. Lapuente-Brun, E. et al. Supercomplex Assembly Determines Electron Flux in the Mitochondrial Electron Transport Chain. Science (1979) 340, 1567–1570 (2013).

60. Zheng, Y., Gibb, A. A., Xu, H., Liu, S. & Hill, B. G. The metabolic state of the heart regulates mitochondrial supercomplex abundance in mice. Redox Biol 63, (2023).

61. Hou, T. et al. NDUFAB1 confers cardio-protection by enhancing mitochondrial bioenergetics through coordination of respiratory complex and supercomplex assembly. Cell Res 29, 754–766 (2019).

62. Rosca, M. G. et al. Cardiac mitochondria in heart failure: Decrease in respirasomes and oxidative phosphorylation. Cardiovasc Res 80, 30–39 (2008).

63. Sehwan Jang et al. Elucidating Mitochondrial Electron Transport Chain Supercomplexes in the Heart During Ischemia– Reperfusion. Antioxid Redox Signal 27, (2017).

64. Stroud, D. A. et al. Accessory subunits are integral for assembly and function of human mitochondrial complex i. Nature 538, 123–126 (2016).

65. Van Vranken, J. G. et al. The mitochondrial acyl carrier protein (ACP) coordinates mitochondrial fatty acid synthesis with iron sulfur cluster biogenesis. (2016) doi:10.7554/eLife.17828.001.

66. Cogliati, S. et al. Mechanism of super-assembly of respiratory complexes III and IV. Nature 539, 579–582 (2016).

67. Lobo-Jarne, T. et al. Human COX7A2L Regulates Complex III Biogenesis and Promotes Supercomplex Organization Remodeling without Affecting Mitochondrial Bioenergetics. Cell Rep 25, 1786–1799.e4 (2018).

68. Benegiamo, G. et al. COX7A2L genetic variants determine cardiorespiratory fitness in mice and human. Nat Metab 4, 1336–1351 (2022).

69. Ragone, I. et al. SERCA2a Protein Levels Are Unaltered in Human Heart Failure. Circulation 148, 613–616 (2023).

70. Hoppins, S. et al. A mitochondrial-focused genetic interaction map reveals a scaffold-like complex required for inner membrane organization in mitochondria. Journal of Cell Biology 195, 323–340 (2011).

71. Barbot, M. et al. Mic10 Oligomerizes to Bend Mitochondrial Inner Membranes at Cristae Junctions. Cell Metab 21, 756–763 (2015).

72. Körner, C. et al. The C-terminal domain of Fcj1 is required for formation of crista junctions and interacts with the TOB/SAM complex in mitochondria. Mol Biol Cell 23, 2143–2155 (2012).

73. Schweppe, D. K. et al. Mitochondrial protein interactome elucidated by chemical cross-linking mass spectrometry. Proc Natl Acad Sci U S A 114, 1732–1737 (2017).

74. Tang, J. et al. Sam50–Mic19–Mic60 axis determines mitochondrial cristae architecture by mediating mitochondrial outer and inner membrane contact. Cell Death Differ 27, 146–160 (2020).

75. Friedman, J. R., Mourier, A., Yamada, J., Mccaffery, J. M. & Nunnari, J. MICOS coordinates with respiratory complexes and lipids to establish mitochondrial inner membrane architecture. doi:10.7554/eLife.07739.001.

76. Anand, R. et al. MIC26 and MIC27 cooperate to regulate cardiolipin levels and the landscape of OXPHOS complexes. Life Sci Alliance 3, (2020).

77. Bohnert, M. et al. Central Role of Mic10 in the Mitochondrial Contact Site and Cristae Organizing System. Cell Metab 21, 747–755 (2015).

78. Tirrell, P. S., Nguyen, K. N., Luby-Phelps, K. & Friedman, J. R. MICOS subcomplexes assemble independently on the mitochondrial inner membrane in proximity to ER contact sites. Journal of Cell Biology 219, (2020).

79. Qiu, J. et al. Coupling of mitochondrial import and export translocases by receptor-mediated supercomplex formation. Cell 154, (2013).

80. Burri, L. et al. Mature DIABLO/Smac Is Produced by the IMP Protease Complex on the Mitochondrial Inner Membrane. Mol Biol Cell 16, 2926–2933 (2005).

81. Nielsen, M. S., van Opbergen, C. J. M., van Veen, T. A. B. & Delmar, M. The intercalated disc: a unique organelle for electromechanical synchrony in cardiomyocytes. Physiol Rev 103, 2271–2319 (2023).

82. Hulsmans, M. et al. Macrophages Facilitate Electrical Conduction in the Heart. Cell 169, 510–522.e20 (2017).

83. Sugita, J. et al. Cardiac macrophages prevent sudden death during heart stress. Nat Commun 12, (2021).

84. Vasquez, C. et al. Enhanced fibroblast-myocyte interactions in response to cardiac injury. Circ Res 107, 1011–1020 (2010).

85. Hall, C., Gehmlich, K., Denning, C. & Pavlovic, D. Complex relationship between cardiac fibroblasts and cardiomyocytes in health and disease. J Am Heart Assoc 10, 1–15 (2021).

86. Yang, Z. et al. Desmosomal dysfunction due to mutations in desmoplakin causes arrhythmogenic right ventricular dysplasia/cardiomyopathy. Circ Res 99, 646–655 (2006).

87. Syrris, P. et al. Arrhythmogenic Right Ventricular Dysplasia/Cardiomyopathy Associated with Mutations in the Desmosomal Gene Desmocollin-2. The American Journal of Human Genetics 79, 978–984 (2006).

88. Norman, M. et al. Novel mutation in desmoplakin causes arrhythmogenic left ventricular cardiomyopathy. Circulation 112, 636–642 (2005).

89. Li, J. et al. Cardiac-specific loss of N-cadherin leads to alteration in connexins with conduction slowing and arrhythmogenesis. Circ Res 97, 474–481 (2005).

90. Ghidoni, A. et al. Cadherin 2-Related Arrhythmogenic Cardiomyopathy: Prevalence and Clinical Features. Circ Genom Precis Med 14, E003097 (2021).

91. Kostetskii, I. et al. Induced deletion of the N-cadherin gene in the heart leads to dissolution of the intercalated disc structure. Circ Res 96, 346–354 (2005).

92. Mayosi, B. M. et al. Identification of Cadherin 2 (CDH2) Mutations in Arrhythmogenic Right Ventricular Cardiomyopathy. Circ Cardiovasc Genet 10, (2017).

93. Lyon, R. C. et al. Connexin defects underlie arrhythmogenic right ventricular cardiomyopathy in a novel mouse model. Hum Mol Genet 23, 1134–1150 (2014).

94. Van Norstrand, D. W. et al. Connexin43 mutation causes heterogeneous gap junction loss and sudden infant death. Circulation 125, 474–481 (2012).

95. Campbell, I. D. & Humphries, M. J. Integrin structure, activation, and interactions. Cold Spring Harb Perspect Biol 3, 1–14 (2011).

96. Dave, J. et al. Gene editing reverses arrhythmia susceptibility in humanized PLN-R14del mice: modelling a European cardiomyopathy with global impact. Cardiovasc Res (2022) doi:10.1093/cvr/cvac021.

97. Haghighi, K. et al. Impaired right ventricular calcium cycling is an early risk factor in r14del-phospholamban arrhythmias. J Pers Med 11, (2021).

98. Peng, T. I. & Jou, M. J. Oxidative stress caused by mitochondrial calcium overload. Ann N Y Acad Sci 1201, 183–188 (2010).

99. Santulli, G., Xie, W., Reiken, S. R. & Marks, A. R. Mitochondrial calcium overload is a key determinant in heart failure. Proc Natl Acad Sci U S A 112, 11389–11394 (2015).

100. Eisner, D. A., Caldwell, J. L., Trafford, A. W. & Hutchings, D. C. The Control of Diastolic Calcium in the Heart: Basic Mechanisms and Functional Implications. Circ Res 126, 395–412 (2020).

101. Kaushik, S. & Cuervo, A. M. Proteostasis and aging. Nat Med 21, 1406–1415 (2015).

102. Henning, R. H. & Brundel, B. J. Proteostasis in cardiac health and disease. Nat Rev Cardiol 14, 637–653 (2017).

103. Sharma, P. et al. Endoplasmic reticulum protein targeting of phospholamban: A common role for an N-terminal di-arginine motif in ER retention? PLoS One 5, (2010).

104. Haghighi, K. et al. The human phospholamban Arg14-deletion mutant localizes to plasma membrane and interacts with the Na/K-ATPase. J Mol Cell Cardiol 52, 773–782 (2012).

105. Michelsen, K., Yuan, H. & Schwappach, B. Hide and run: Arginine-based endoplasmic-reticulum-sorting motifs in the assembly of heteromultimeric membrane proteins. EMBO Rep 6, 717–722 (2005).

106. Arakel, E. C. et al. Tuning the electrical properties of the heart by differential trafficking of KATP ion channel complexes. J Cell Sci 127, 2106–2119 (2014).

107. Yaffe, M. B. et al. The Structural Basis for 14-3-3:Phosphopeptide Binding Specificity. Cell 91, 961–971 (1997).

108. Smith, A. J., Daut, J. & Schwappach, B. Membrane proteins as 14-3-3 clients in functional regulation and intracellular transport. Physiology 26, 181–191 (2011).

109. Nielsen, M. S., van Opbergen, C. J. M., van Veen, T. A. B. & Delmar, M. The intercalated disc: a unique organelle for electromechanical synchrony in cardiomyocytes. Physiol Rev 103, 2271–2319 (2023).

110. Moise, N., Struckman, H. L., Dagher, C., Veeraraghavan, R. & Weinberg, S. H. Intercalated disk nanoscale structure regulates cardiac conduction. Journal of General Physiology 153, (2021).

111. Kucera, J. P., Rohr, S. & Rudy, Y. Localization of sodium channels in intercalated disks modulates cardiac conduction. Circ Res 91, 1176–1182 (2002).

112. Asimaki, A. et al. A novel dominant mutation in plakoglobin causes arrhythmogenic right ventricular cardiomyopathy. Am J Hum Genet 81, 964–973 (2007).

113. Rizzo, S. et al. Intercalated disc abnormalities, reduced Na+ current density, and conduction slowing in desmoglein-2 mutant mice prior to cardiomyopathic changes. Cardiovasc Res 95, 409–418 (2012).

114. Ito, Y. et al. Disorganization of intercalated discs in dilated cardiomyopathy. Sci Rep 11, (2021).

115. Kudo-Sakamoto, Y. et al. Calpain-dependent cleavage of N-cadherin is involved in the progression of post-myocardial infarction remodeling. Journal of Biological Chemistry 289, 19408–19419 (2014).

116. Brundel, B. J. J. M., et al. Activation of Proteolysis by Calpains and Structural Changes in Human Paroxysmal and Persistent Atrial Fibrillation. Cardiovascular Research vol. 54 www.elsevier.com/locate/cardiores (2002).

117. Weninger, G. et al. Calpain cleavage of Junctophilin-2 generates a spectrum of calcium-dependent cleavage products and DNA-rich NT1-fragment domains in cardiomyocytes. Sci Rep 12, (2022).

118. Wu, C. Y. C. et al. Calpain-dependent cleavage of junctophilin-2 and T-tubule remodeling in a mouse model of reversible heart failure. J Am Heart Assoc 3, (2014).

119. Weninger, G. & Lehnart, S. E. New Insights into the Proteolytic Regulation of the Structural Protein Junctophilin-2 by Calpain. J Cell Signal 3, 171–178 (2022).

120. Qi, B. et al. Cardiac-specific overexpression of Ndufs1 ameliorates cardiac dysfunction after myocardial infarction by alleviating mitochondrial dysfunction and apoptosis. Exp Mol Med 54, 946–960 (2022).

121. Graham, D. et al. Mitochondria-targeted antioxidant mitoq10 improves endothelial function and attenuates cardiac hypertrophy. Hypertension 54, 322–328 (2009).

122. Smith, R. A. J. & Murphy, M. P. Animal and human studies with the mitochondria-targeted antioxidant MitoQ. Ann N Y Acad Sci 1201, 96–103 (2010).

123. Daubert, M. A. et al. Novel Mitochondria-Targeting Peptide in Heart Failure Treatment: A Randomized, Placebo-Controlled Trial of Elamipretide. Circ Heart Fail 10, (2017).

124. Compagnoni, G. M. et al. The Role of Mitochondria in Neurodegenerative Diseases: the Lesson from Alzheimer’s Disease and Parkinson’s Disease. Mol Neurobiol 7, 2959–2980 (2020).

125. Ribeiro Junior, R. F., et al. MitoQ improves mitochondrial dysfunction in heart failure induced by pressure overload. Free Radic Biol Med 117, 18–29 (2018).

126. 126. Souza-Neto, F. V., et al. Mitochondrial Oxidative Stress Promotes Cardiac Remodeling in Myocardial Infarction through the Activation of Endoplasmic Reticulum Stress. Antioxidants 11, (2022).

127. Snow, B. J. et al. A double-blind, placebo-controlled study to assess the mitochondria-targeted antioxidant MitoQ as a disease-modifying therapy in Parkinson’s disease. Movement Disorders 25, 1670–1674 (2010).

128. Gane, E. J. et al. The mitochondria-targeted anti-oxidant mitoquinone decreases liver damage in a phase II study of hepatitis C patients. Liver International 30, 1019–1026 (2010).

129. Cho, J. et al. Potent mitochondria-targeted peptides reduce myocardial infarction in rats. Coron Artery Dis 18, 215–220 (2007).

130. Linke, D. Chapter 34 Detergents. An Overview. in Methods in Enzymology vol. 463 603–617 (Academic Press Inc., 2009).

131. Frewen, B. E., Merrihew, G. E., Wu, C. C., Noble, W. S. & MacCoss, M. J. Analysis of peptide MS/MS spectra from large-scale proteomics experiments using spectrum libraries. Anal Chem 78, 5678–5684 (2006).

132. Neef, J. et al. Quantitative optical nanophysiology of Ca2+ signaling at inner hair cell active zones. Nat Commun 9, (2018).

133. Wagner, E., Brandenburg, S., Wagner, E. & Lehnart, S. E. Analysis of tubular membrane networks in cardiac myocytes from atria and ventricles. Journal of Visualized Experiments (2014) doi:10.3791/51823.

134. Atanassov, I. & Urlaub, H. Increased proteome coverage by combining PAGE and peptide isoelectric focusing: Comparative study of gel-based separation approaches. Proteomics 13, 2947–2955 (2013).

135. Meier, F. et al. Parallel accumulation-serial fragmentation (PASEF): Multiplying sequencing speed and sensitivity by synchronized scans in a trapped ion mobility device. J Proteome Res 14, 5378–5387 (2015).

136. Meier, F. et al. Online parallel accumulation–serial fragmentation (PASEF) with a novel trapped ion mobility mass spectrometer. Molecular and Cellular Proteomics 17, 2534–2545 (2018).

137. Skowronek, P. et al. Rapid and In-Depth Coverage of the (Phospho-) Proteome With Deep Libraries and Optimal Window Design for dia-PASEF. Molecular and Cellular Proteomics 21, (2022).

138. Wittig, I., Braun, H. P. & Schägger, H. Blue native PAGE. Nat Protoc 1, 418–428 (2006).

139. Pearson, K. & Lee, A. On the laws of inheritance in man: I. Inheritance of physical characters. Biometrika 2, 357–462 (1903).

140. Bateman, A. et al. UniProt: the Universal Protein Knowledgebase in 2023. Nucleic Acids Res 51, D523–D531 (2023).

141. Mi, H., Muruganujan, A., Ebert, D., Huang, X. & Thomas, P. D. PANTHER version 14: More genomes, a new PANTHER GO-slim and improvements in enrichment analysis tools. Nucleic Acids Res 47, D419–D426 (2019).

142. Thomas, P. D., et al. PANTHER: Making genome-scale phylogenetics accessible to all. Protein Science vol. 31 8–22 Preprint at 10.1002/pro.4218 (2022).

143. Mi, H., Muruganujan, A., Casagrande, J. T. & Thomas, P. D. Large-scale gene function analysis with the panther classification system. Nat Protoc 8, 1551–1566 (2013).

144. Perez-Riverol, Y. et al. The PRIDE database resources in 2022: A hub for mass spectrometry-based proteomics evidences. Nucleic Acids Res 50, D543–D552 (2022).

145. Doevendans, P. A., Glijnis, P. C. & Kranias, E. G. Leducq Transatlantic Network of Excellence to Cure Phospholamban-Induced Cardiomyopathy (CURE-PLaN). Circulation Research vol. 125 720–724 Preprint at 10.1161/CIRCRESAHA.119.315077 (2019).

