## Supplemental figures for "Unbiased complexome profiling and global proteomics analysis reveals mitochondrial impairment and potential changes at the intercalated disk in presymptomatic R14^Δ/+^ mice hearts"

### Supplemental Figures and Tables

A)

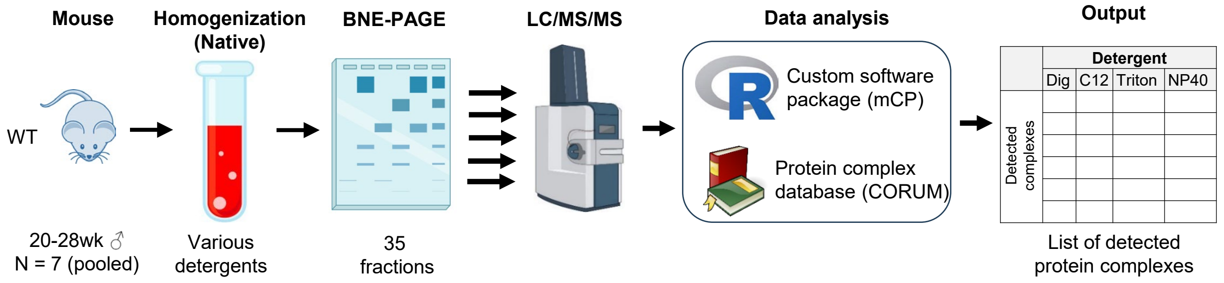


B)

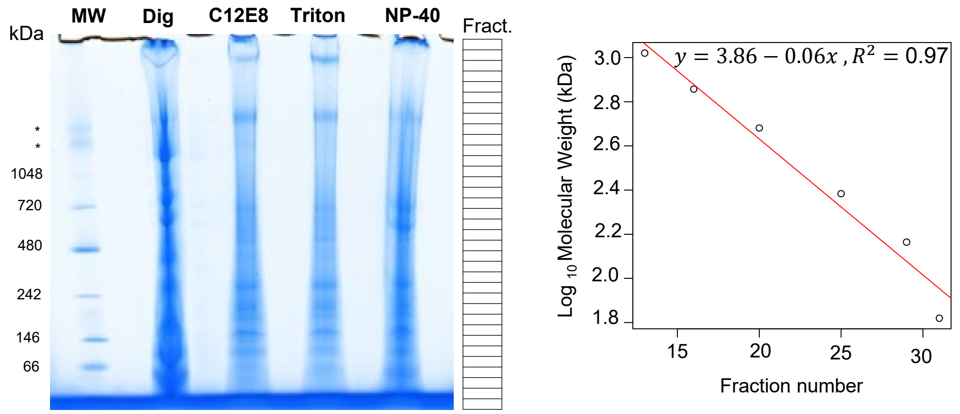


C)

| Protein complexes | Digitonin | C12E8 | Triton | NP-40 |
|---|---|---|---|---|
| Protein complexes detected | 26 | 32 | 30 | 30 |
| Unique complexes | 1 | 1 | 3 | 1 |
| Shared with all detergents | 21 | 21 | 21 | 21 |
| Shared with ≥1 other detergent | 4 | 10 | 6 | 8 |

**Supplemental S1**: Optimization of detergent conditions for cardiac tissue samples.
**A)** Detergent optimization workflow. Enriched membrane fractions were prepared from left-ventricular tissues extracted from 21-28wk-old ♂ mice (N = 7). Membrane preparations were pooled, then solubilized using either digitonin (6mg/mg protein), C12E8 (0.5mg/mg protein), Triton X-100 (0.01mg/mg protein) or NP-40 (0.5mg/mg protein)/ Solubilized membranes were fractionated into 35 fractions via BN-PAGE and subject to MS-based proteomics analysis. A custom R-script (mCP, manuscript in preparation) was used to identify protein complexes within the resultant dataset, using the CORUM v. 4.0 protein complex library as a ground-truth. **B)** BN-PAGE analysis of ventricular membrane preparations solubilized in digitonin, C12E8, Triton X-100 or NP40. Approximate fractions used for complexome-profiling analysis shown to the left. Molecular-weight calibration shown on the far left. **C)** Number of protein complexes detected following solubilization with the listed detergent. Use of C12E8 resulted in detection of a greater number of proteins. Full list of protein-complex IDs in Supplemental S2. Protein complexes detected using a custom R-script coupled to the CORUM protein complex library as described in A).

| CORUM protein complex detected in C12E8 solubilization | Detected subunits | Total subunits |
|---|---|---|
| IDH3G STRING COMPLEX | 10 | 10 |
| Caveolins and others | 6 | 14 |
| 20S proteasome | 14 | 14 |
| Immunoproteasome | 13 | 14 |
| COP9 signalosome complex | 4 | 8 |
| CCT complex (chaperonin containing TCP1 complex)_1 | 8 | 24 |
| CCT complex (chaperonin containing TCP1 complex)_2 | 8 | 24 |
| Skeletal muscle sarcoglycan complex SGC, alpha-beta-gamma-delta | 4 | 12 |
| CCT complex (chaperonin containing TCP1 complex)_3 | 8 | 24 |
| Skeletal muscle sarcoglycan complex SGC, beta-gamma-delta-zeta | 3 | 4 |
| Smooth muscle sarcoglycan complex SGC, beta-delta-zeta | 2 | 3 |
| Skeletal muscle sarcoglycan complex SGC, alpha-beta-gamma-delta | 4 | 12 |
| Skeletal muscle sarcoglycan complex SGC, alpha-beta-epsilon-gamma | 3 | 4 |
| Skeletal muscle sarcoglycan complex SGC, alpha-beta-gamma-delta | 4 | 12 |
| Skeletal muscle sarcoglycan complex SGC, epsilon-beta-gamma-delta | 3 | 4 |
| Dystrophin-sarcoglycan-syntrophin complex, skeletal muscle | 5 | 6 |
| Sarcoglycan-sarcospan-dystroglycan complex | 5 | 6 |
| Sarcoglycan-sarcospan-complex SG-SPN | 4 | 5 |
| Sarcoglycan-sarcospan-syntrophin-dystrobrevin complex | 4 | 8 |
| Respiratory chain complex I, mitochondrial | 29 | 77 |
| Respiratory chain complex I, mitochondrial | 29 | 77 |
| (ER)-localized multiprotein complex, Ig heavy chains associated | 7 | 10 |
| Succinate dehydrogenase complex II, mitochondrial | 4 | 4 |
| Cytochrome bc1-complex, mitochondrial | 9 | 10 |
| Cytochrome c oxidase, mitochondrial | 11 | 13 |
| Kif13a-AP1 complex | 2 | 4 |
| Itgav-Itgb3-Gsn complex | 3 | 3 |
| Parvulin-associated pre-rRNP complex | 23 | 62 |
| CCT complex (chaperonin containing TCP1 complex), testis specific | 7 | 8 |
| Vps29-Vps35-Vps26a complex | 2 | 3 |
| Dnajc5-Sgta complex | 2 | 3 |
| Ap1g1-Ap2a2-Dcx complex | 2 | 3 |

**Supplemental S2**: List of protein complexes detected in Supplemental S1C

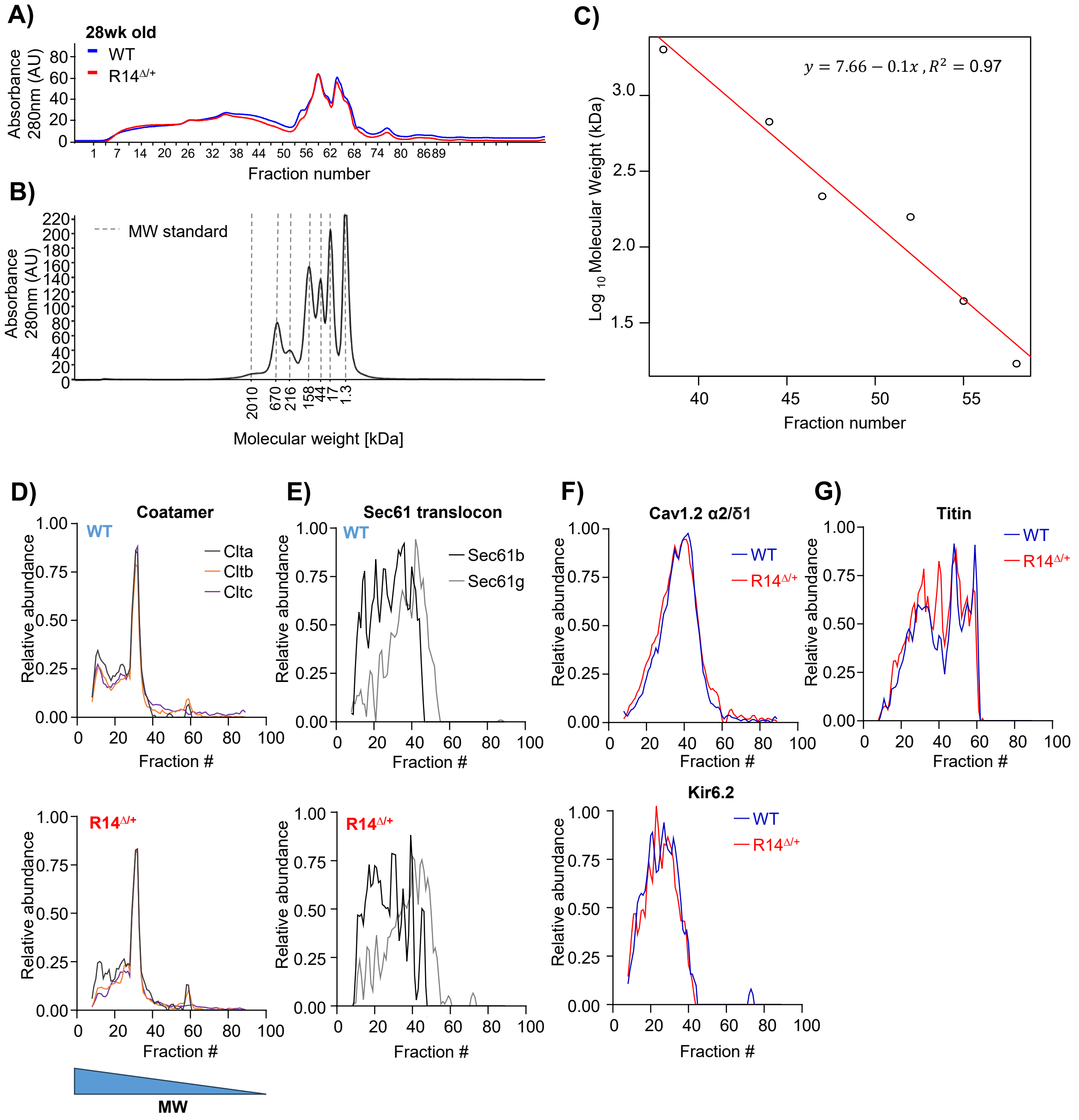


**Supplemental S3**: SEC-MS analysis of 28wk-old adult WT and $R14^{\Delta/+}$ mice

**A-C)** UV-absorption chromatogram (A) and molecular-weight calibration curve (B-C) for SEC-MS experiment shown in Figure 1. **D-G)** Elution profiles for additional protein complexes located in cytosolic (Coatamer), integral SER membrane (Sec61-translocon), integral plasma-membrane (Cav1.2 and Kir6.2 multimers) and sarcomeric (Titin) compartments. Elution profiles subject to curve smoothing as in Figure 1b-e.

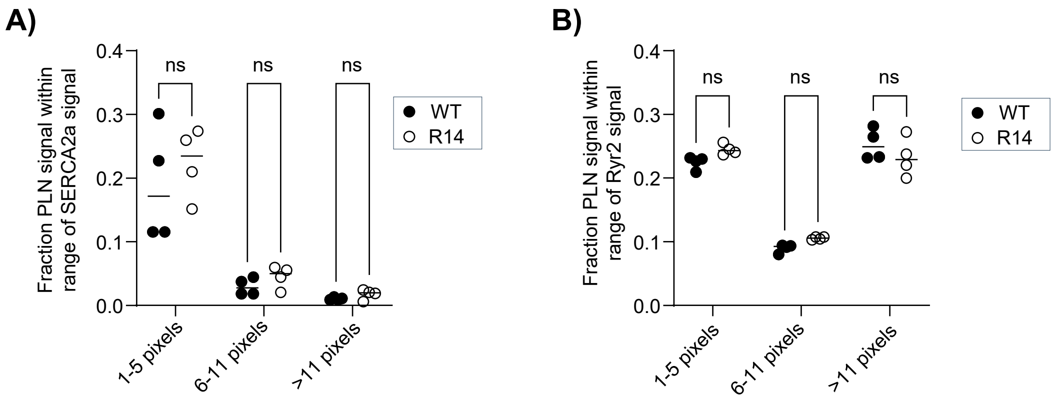


**Supplemental S4**: Additional quantification of PLN-SERCA2a and PLN-RyR2 interactions in adult 28wk-old mice. **A-B)** Fraction of PLN pixels within a given distance from nearest SERCA2a (A) or RyR2 (B) signal. Representative immunofluorescence images shown in Figure 2. Statistical significance determined via two-way ANOVA with Šidák correction for multiple comparisons (ns: no significant difference).

A)

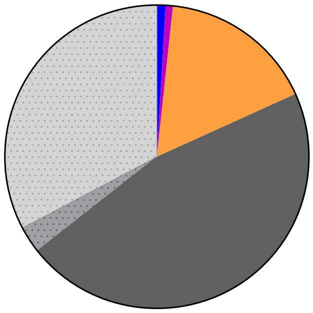


B)

| Complex | | Source | |
| --- | --- | --- | --- |
| ID# | Name | Organism | Material |
| 6752 | Smad3-mSin3A-Hdac1 complex | Mouse | cardiac fibroblasts |
| 6241 | PMCA4-alpha-1-syntrophin-NOS-1 complex | Mouse | cardiac muscle |
| 6240 | PMCA1-alpha-1-syntrophin-NOS-1 complex | Mouse | cardiac muscle cells |
| 6271 | CASQ2-RYR1-ASPH-TRDN complex | Dog | cardiac muscle cells |
| 6774 | Fhl1-Pdlim1-Gsn-Actn1 complex | Mouse | cardiac ventricle lysates |
| 6672 | Ank2-Kcnj11-Abcc8 complex | Mouse | cardiomyocites |
| 6673 | Foxo1-Yap1 complex | Mouse | cardiomyocytes |
| 6674 | Ank2-Kcnj11-Abcc9 complex | Mouse | cardiomyocytes |
| 7003 | Cacna1d-Kcnn4 complex | Mouse | cardiomyocytes |
| 6654 | KCNJ11-ABCC9 complex | Human | COS cells, cardiomyocytes |
| 5493 | Tmsb4x-Lims1-Ilk complex | Mouse | COS, 10T1/2 cells, heart muscle |
| 1071 | PKD2-FPC complex | Human | epithelium, kidney, heart muscle |
| 178 | Respiratory chain complex I (holoenzyme), mitochondrial | Human | heart |
| 6879 | Dystrophin-glycoprotein complex (DGC) | Mouse | heart |
| 7475 | Wdr26-Vdac1 complex | Rat | heart |
| 7481 | Nos1-Xdh complex | Mouse | heart |
| 7569 | Ldb3(CypherS)-Myoz2-Pdlim5 complex | Mouse | heart |
| 8036 | Fhl1-Titin complex | Mouse | heart |
| 8037 | Fhl1-Titin-Raf1-Mek2-Erk2 complex | Mouse | heart |
| 384 | Caveolar macromolecular signaling complex | Mouse | heart muscle |
| 385 | Caveolar macromolecular signaling complex, anti-Cav-3 | Mouse | heart muscle |
| 399 | Respiratory chain complex I, mitochondrial | Bovine | heart muscle |
| 400 | Respiratory chain complex I (subcomplex I alpha) | Bovine | heart muscle |
| 401 | Respiratory chain complex I (subcomplex I lambda) | Bovine | heart muscle |
| 402 | Respiratory chain complex I (subcomplex I beta) | Bovine | heart muscle |
| 403 | Cytochrome bc1-complex (EC 1.10.2.2), mitochondrial | Bovine | heart muscle |
| 421 | Dlg1-Cav3-Kcna5 complex | Rat | heart muscle |
| 520 | KCNQ1 macromolecular complex | Human | heart muscle |
| 563 | F1F0-ATPase, mitochondrial | Human | heart muscle |
| 574 | F1-ATPase-IF1 (inhibitor protein) complex, mitochondrial | Bovine | heart muscle |
| 576 | F1F0-ATPase, mitochondrial | Bovine | heart muscle |
| 584 | Ppp3cb-RyR2-mAkap complex | Rat | heart muscle |
| 589 | Respiratory chain supercomplex (complex I, III, IV), mito., heart | Bovine | heart muscle |
| 630 | RyR2 macromolecular complex | Rat | heart muscle |
| 737 | Ppp2c-PKA-Ryr2-mAkap complex | Rat | heart muscle |
| 779 | K(ATP) macromolecular complex (Kir6.2, Pkm2, Gapdh, Tpi1) | Rat | heart muscle |
| 2689 | Fra1-JunB DNA-protein complex | Rat | heart muscle |
| 2796 | Fosb-Junb DNA-protein complex | Rat | heart muscle |
| 2953 | Respiratory chain complex I (nuclear encoded subunits) | Bovine | heart muscle |
| 3867 | Xin-Cdh2-Ctnnb1-Ctnnd1 complex | Mouse | heart muscle |
| 8295 | Smarca4-Foxm1 complex | Mouse | heart ventricle |
| 6865 | Smarca4-Ehmt2-Dnmt3a complex | Mouse | Hearts from E12.5 mouse embryos and adult mice |
| 6510 | ILK-SERCA2A-PLN complex | Human | iPSC-derived cardiomyocytes |
| 6711 | Calpain 2 | Bovine | myocardial cells |
| 7927 | AGTR1-AGTR2 complex | Human | myometrium |
| 7503 | Potassium channel complex (Kcnb1, Kcne2, Kcne1, Kcnh2) | Rat | rat heart |
| 7511 | Kcnb1-Kcne1 complex | Rat | rat heart tissue |
| 7512 | Kcnb1-Kcne2 complex | Rat | rat heart tissue |

**Supplemental S5**: Suitability of CORUM v. 4.0 for analysis of cardiac samples
**A)** Origins of protein complexes documented in CORUM v. 4.0. Database entries were manually annotated based on given source. **B)** Complete list of CORUM entries observed in cardiac-related sources.

| **CORUM protein complex detected in 28wk-old ♂ WT and R14$^{\Delta/+}$ mice** | **Detected subunits** | **Total subunits** |
|---|---|---|
| IDH3G STRING COMPLEX | 6 | 8 |
| 9S-cytosolic aryl hydrocarbon (Ah) receptor non-ligand activated complex | 10 | 10 |
| 20S proteasome | 3 | 4 |
| Immunoproteasome | 14 | 14 |
| COP9 signalosome complex | 13 | 14 |
| CCT complex (chaperonin containing TCP1 complex) | 6 | 8 |
| CCT complex (chaperonin containing TCP1 complex) | 8 | 24 |
| Gamma-secretase complex (Aph1a, Psen1, Psenen, Ncstn) | 8 | 24 |
| Skeletal muscle sarcoglycan complex SGC, alpha-beta-gamma-delta | 2 | 8 |
| CCT complex (chaperonin containing TCP1 complex) | 4 | 12 |
| Skeletal muscle sarcoglycan complex SGC, beta-gamma-delta-zeta | 8 | 24 |
| Skeletal muscle sarcoglycan complex SGC, alpha-beta-gamma-delta | 3 | 4 |
| Skeletal muscle sarcoglycan complex SGC, alpha-beta-epsilon-gamma | 4 | 12 |
| Skeletal muscle sarcoglycan complex SGC, alpha-beta-gamma-delta | 3 | 4 |
| Skeletal muscle sarcoglycan complex SGC, epsilon-beta-gamma-delta | 4 | 12 |
| Dystrophin-sarcoglycan-syntrophin complex, skeletal muscle | 3 | 4 |
| Sarcoglycan-sarcospan-dystroglycan complex | 5 | 6 |
| Sarcoglycan-sarcospan-complex SG-SPN | 5 | 6 |
| Sarcoglycan-sarcospan-syntrophin-dystrobrevin complex | 4 | 5 |
| Respiratory chain complex I, mitochondrial | 5 | 8 |
| Respiratory chain complex I, mitochondrial | 30 | 77 |
| Succinyl-CoA synthetase, ADP-forming | 30 | 77 |
| Succinyl-CoA synthetase, GDP-forming | 2 | 2 |
| (ER)-localized multiprotein complex, Ig heavy chains associated | 2 | 2 |
| Succinate dehydrogenase complex II, mitochondrial | 8 | 10 |
| Cytochrome bc1-complex, mitochondrial | 4 | 4 |
| Cytochrome c oxidase, mitochondrial | 9 | 10 |
| Gamma-secretase complex (Aph1a, Ncstn, Psen1, Psenen, Tmp21) | 10 | 13 |
| TRPV5-S100A10-annexin 2 complex | 3 | 5 |
| S100A10-annexin 2 complex | 2 | 3 |
| SNARE complex (Vti1b, Stx6, Stx7) | 2 | 2 |
| SNARE complex (Vti1b, Stx7, Stx8, Vamp8) | 2 | 3 |
| SNARE complex (Stx4, Stx6, Stx7, Vamp3, Vamp7, Vamp8, Vti1b) | 3 | 4 |
| Kif13a-AP1 complex | 5 | 7 |
| ERdj3-BiP complex | 3 | 4 |
| Itgav-Itgb3-Gsn complex | 2 | 2 |
| Itga-Itgb1-Ppap2b complex | 2 | 3 |
| Aph1a-Psen1-Ncstn complex | 3 | 3 |
| Gata1-Fog1-MeCP1 complex | 2 | 3 |
| Nkx3.2-SMAD1-SMAD4-HDAC-Sin3A complex | 3 | 13 |
| Drosha complex | 2 | 7 |
| Parvulin-associated pre-rRNP complex | 5 | 8 |
| CCT complex (chaperonin containing TCP1 complex), testis specific | 27 | 62 |
| G protein complex (Hdac4, Gnb1, Gng2) | 7 | 8 |
| G protein complex (Hdac5, Gnb1, Gng2) | 2 | 3 |
| G protein complex (Btk, Gng2, Gnb1) | 2 | 3 |
| Vps29-Vps35-Vps26a complex | 2 | 3 |
| Gamma-secretase complex (Aph1a, Psen1, Psenen, Ncstn) | 2 | 3 |
| Ksr1-PP2A holoenzyme complex (Ppp2r1a, Ppp2r2b, Ppp2ca), PDGF stimulated | 2 | 8 |
| Ksr1-PP2A core enzyme complex (Ppp2r1a, Ppp2ca), untreated | 2 | 4 |
| Ksr1-PP2A core enzyme complex (Ppp2r1a, Ppp2ca), untreated | 2 | 6 |
| Raf1-PP2A holoenzyme complex (Ppp2r1a, Ppp2r2b, Ppp2ca), PDGF stimulated | 2 | 6 |
| Raf1-PP2A core enzyme complex (Ppp2r1a, Ppp2ca), untreated | 3 | 4 |
| p18-p14-Mp1 complex | 3 | 3 |
| Mp1-p14 scaffolding complex | 3 | 3 |
| Ncstn-Psen1 complex | 2 | 2 |
| Rabep1-Ap1g1-Ap1s1 complex | 2 | 2 |

**Supplemental S6**: List of CORUM-defined protein complexes detected in CP analysis of adult 28wk-old WT and R14$^{\Delta/+}$ mice. Data was analyzed using the mCP custom R-script as described in Supplemental figure S1.

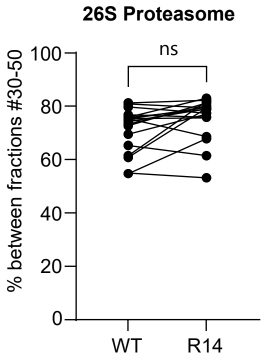


**Supplemental S7**: 26S Proteasome is unchanged in adult 28wk R14$^{\Delta/+}$ mice. The area under the curve of each protein was calculated as in Figure 5. Elution profiles shown in Figure 1D. Significance determined via mixed one-way ANOVA with Šidák corrections for multiple comparison (ns: no significant difference).

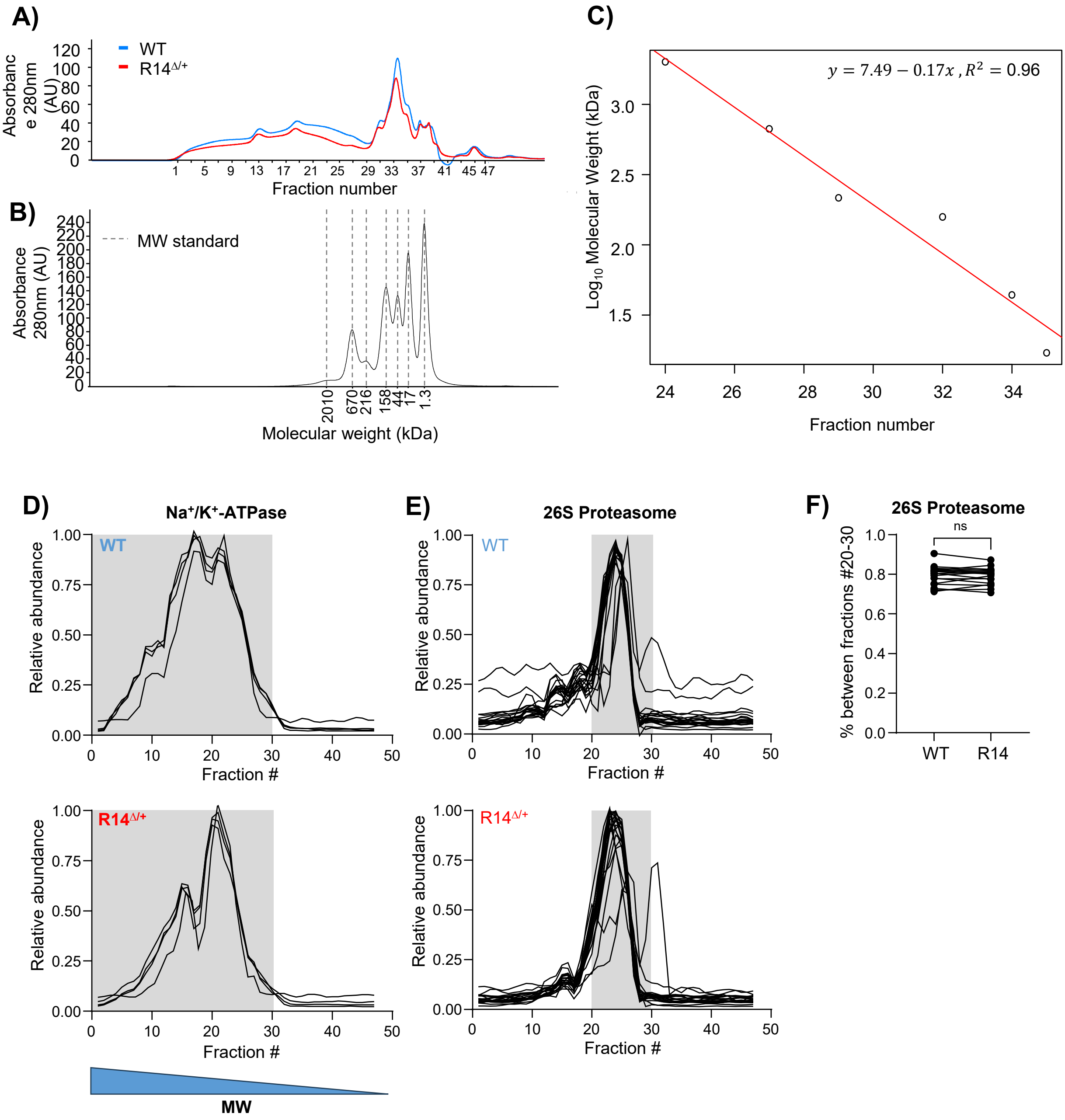


**Supplemental S8**: SEC-MS analysis of 9wk-old juvenile WT and R14$^{\Delta/+}$ mice

**A-C)** UV-absorption chromatogram (A) and molecular-weight calibration curve (B-C) for SEC-MS experiment shown in Figure 7. **D-E)** Elution profiles for $Na^+/K^+$-ATPase and 26S proteasome subunits. Each trace represents the profile of a single protein. Full list of protein IDs in Table 4.**F)** Quantification of supercomplex integrity. Quantification based on the proportion of the area under the curve within the defined peaks as in Figure 7. Significance determined via mixed one-way ANOVA with Šidák corrections for multiple comparison (ns: no significant difference). Elution profiles subject to curve smoothing as in Figure 1b-e.

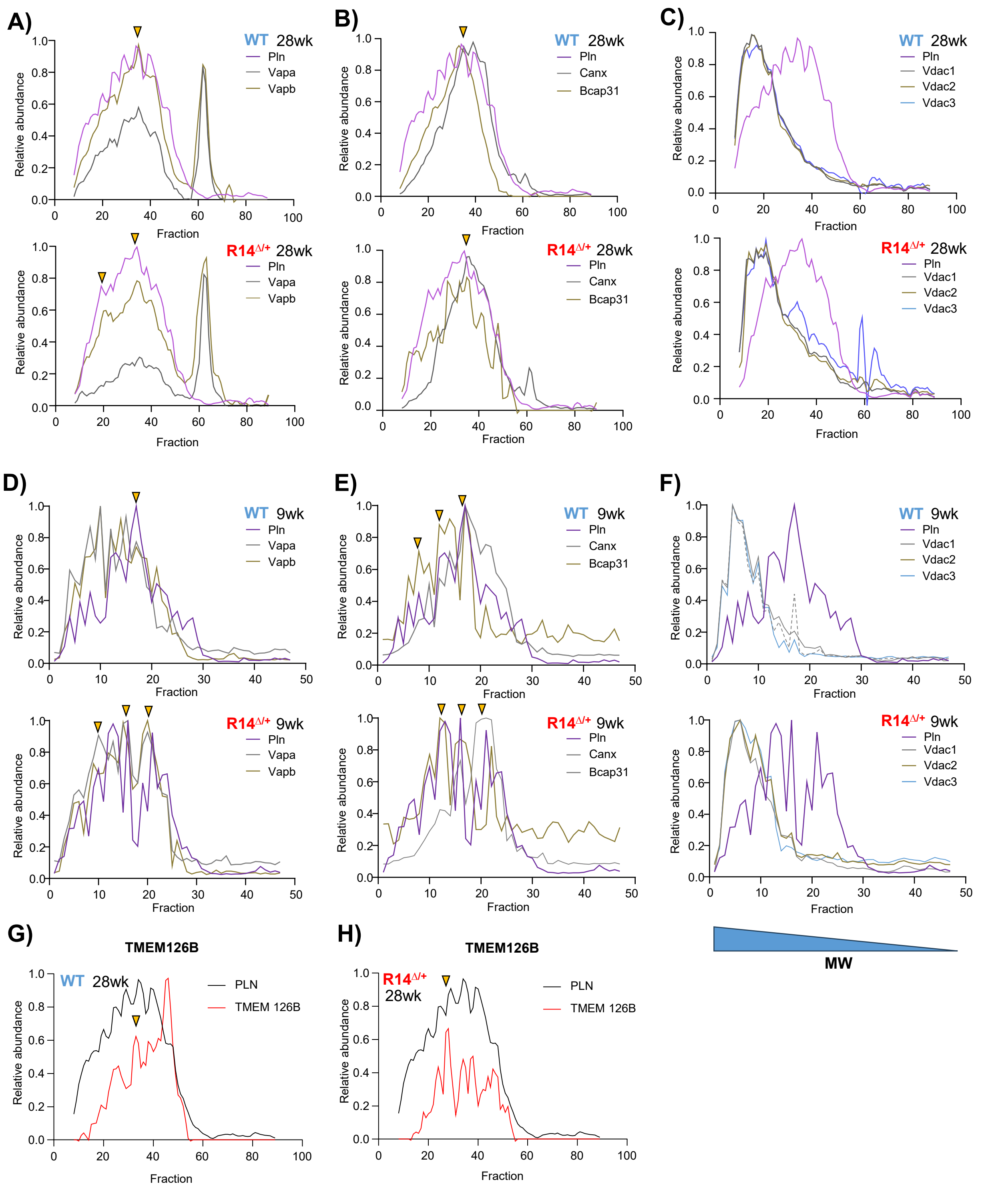


**Supplemental S9**: PLN co-elutes with a subset of ER-mitochondrial tethering proteins and TMEM126B

**A-C)** Elution profiles of PLN (purple) and ER-mitochondrial contact site components VAPA/B, Calnexin (canx), Bcap31 and VDAC1/2/3 from adult 28wk-old WT (top) and R14$^{\Delta/+}$ (bottom) mice. **D-F)** Elution profiles of PLN (purple) and ER-mitochondrial contact site components VAPA/B, Calnexin (canx), Bcap31 and VDAC1/2/3 from juvenile 9wk-old WT (top) and R14$^{\Delta}$/+ (bottom) mice. **G, H)** Coelution of PLN and TMEm126B in 28wk-old WT R14$^{\Delta/+}$ mice (TMEM126B not detected in 9wk-old mouse dataset). Co-fractionating peaks indicated with arrow. Elution profiles subject to curve smoothing as in Figure 1b-e.
